# Locus-specific Convergent Evolution and Interchromosomal Rearrangements Contribute Type I Interferon Diversification in Amniotes

**DOI:** 10.1101/2025.05.08.652797

**Authors:** Le Zhang, Fubo Ma, Jinpeng Liu, Yangchao Yu, Junxiao Ma, Lei Zhang, Bing Li, Chaofan Li, Peng Liu, Liguo Zhang

## Abstract

Type I interferons (IFN-Is) play essential roles in antiviral immune responses. The extensive diversification of IFN-Is into multiple subtypes and redundant gene copies has posed significant challenges for evolutionary reconstruction. To address this, we developed an integrated analytical pipeline combining the IFN-SCOPE classification model and GENE-GRADE algorithm. Through comprehensive synteny-guided phylogenetic analysis, we identified three evolutionarily conserved IFN-I loci (HACD4, MOB3B, and UBAP2) that maintain chromosomal colocalization across all major amniote lineages - mammals, squamates, and Archelosauria (turtles, crocodilians, and birds). While MOB3B-locus maintains a single IFN-κ ortholog, the HACD4 (IFN-H) and UBAP2 (IFN-U) loci show lineage-specific expansion patterns: IFN-H proliferated in mammals/reptiles but remained single-copy in birds, whereas IFN-U expanded in birds but not in other lineages. Phylogenetic analysis reveals these independently evolved multicopy genes nevertheless cluster into two conserved subgroups (IFN-H2/H1 and IFN-U2/U1), suggesting convergent functional specialization. Within IFN-H and IFN-U clusters, the single-copy IFN-H2 and IFN-U2 genes - positioned at the ancestral ends of their respective genomic arrays - likely represent the progenitor sequences of each locus. Notably, among mammalian IFN-H2 genes, we identified the poorly characterized IFN-υ (rather than IFN-β) as the ancestral form of mammalian IFN-H subtypes. Furthermore, we identified IFN-I genes at non-canonical loci resulting from interchromosomal duplication events in tortoises and diving ducks and provide clear evidence that interchromosomal duplications contributed to IFN-I gene diversity. These discoveries advance our understanding of the evolutionary mechanisms shaping intronless IFN-I genes in amniotes, and potentially beneficial for developing novel IFN-I based antiviral treatments through comparative immunological approaches.

## Introduction

Type I interferons (IFN-Is) are a family of cytokines critical for antiviral immune responses ^1,2^. Due to their antiviral properties, IFN-Is have been developed into therapeutic agents for treating viral infections in both human and veterinary medicine ^3,4^. However, prolonged IFN-I signaling is pathogenic and drives autoimmune diseases like systemic lupus erythematosus (SLE). Consequently, therapeutic strategies targeting IFN-I blockade have been developed for SLE treatment^5^.

Based on sequence homology, IFN-Is are classified into multiple subtypes. In humans, the IFN-I family comprises IFN-α (with 13 subtypes), IFN-β, IFN-ω, IFN-ε, and IFN-κ, each exhibiting functional diversity due to differences in receptor affinity, induction pathways, or tissue specificity^3,6,7^. Functional insights into IFN-I subtypes are primarily derived from studies in humans and mice, with limited data from pet and farm animals, and virtually none from other species. Comparative immunology of IFN-I orthologs across diverse taxa could significantly advance our understanding of their evolution and functional diversification. However, identifying these orthologs remains challenging due to the rapid divergence and low sequence conservation of IFN-Is among vertebrates^8–10^.

IFN-I genes originated in fish and have diversified extensively across vertebrate lineages^9,10^. While fish IFN-I genes contain introns, amphibians possess both intron-containing and intronless variants. In contrast, all amniote IFN-Is are intronless, likely arising from a single retroposition event^10–13^.

Here, we present an integrated bioinformatics pipeline combining the IFN-I sequence composition and structure (IFN-SCOPE) classification model and gene-network graph degree centrality (GENE-GRADE algorithm) to comprehensively identify IFN-I coding sequences and analyze their genomic context. Using a synteny-guided phylogenetic approach, we systematically investigate the evolution of IFN-Is in amniotes.

## Materials & Methods

Research methods in this paper can be divided into four sections: unnannotated IFN-I computation, IFN-I conserved loci computation and assignment, IFN-I comparative genomics evolutionary analysis, and, IFN-I functional experiment (**Figure 1**).

**Figure 1.**
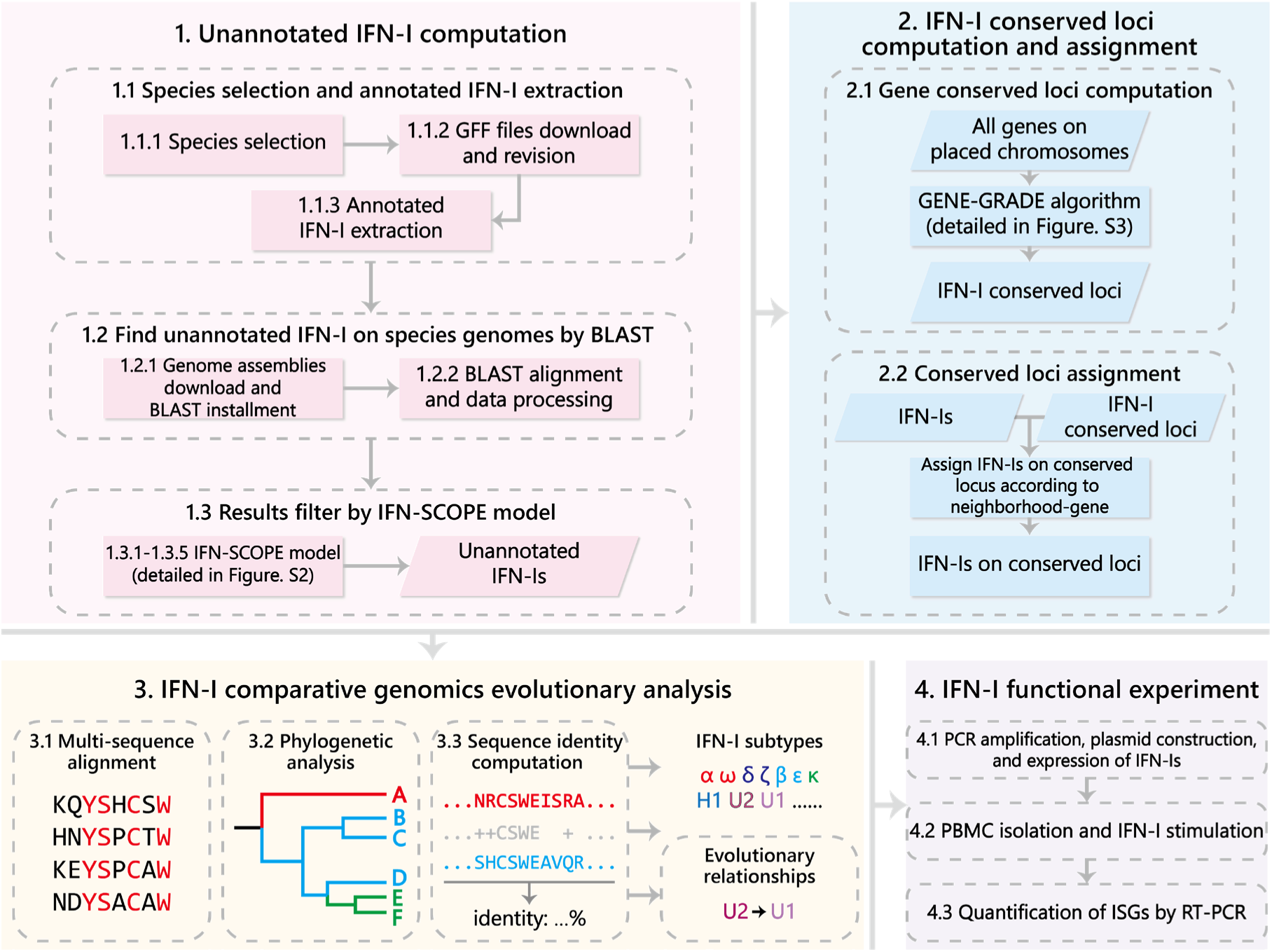
The flowchart of the study.

The numerical labels presented to each segment in the Figure are corresponding to the numerical labels in the Materials & Methods section.

### 1. Unannotated IFN-I computation

#### 1.1 Species selection and annotated IFN-I extraction

##### 1.1.1 Species selection

We first colllect all the order of amniotes from the NCBI taxonomy database. Then, for those orders which have available RefSeq annotation files, we select representative species considering the following principles: species with recently updated RefSeq record, experimental animals, and poultry or livestocks. Next, the phylogenetic relationships of our candidate species are presented (**Figure S1)**.

##### 1.1.2 GFF files download and revision

After determining the species of interest for our study, we download all the general feature format (gff) files (**Table S1**) of these species from the NCBI Assembly database. Then, all the gene names in gff files were comprehensively revised according to their “product”, “description” and “gene abbreviation” terms, which aims to reduce the problem from the non-standardization within the database.

##### 1.1.3 Annotated IFN-I extraction

We filter and extract all annotated IFN-I information through the following steps: 1) include genes containing the fields “ type I” and “interferon/IFN”; 2) Based on step 1), exclude genes containing the fields “type II”, “type III” or “factor”, “induced/inducible” and “stimulated/stimulator”. IFN-I “pseudo genes” and “low-quality proteins” are also excluded in our study **(Table S1)**.

#### 1.2 Find unannotated IFN-I on species genomes by BLAST

To find unannotated IFN-I that are missed by NCBI Eukaryotic Genome Annotation Pipeline, we use BLAST^14^ to find possible protein-coding regions of IFN-I on the genomes of candidate species.

##### 1.2.1 Genome assemblies download and BLAST installment

We download genome assemblies of our candidate species from NCBI Assembly database

**(Table S1)**, and install a linux version of BLAST+ (v2.13.0).

##### 1.2.2 BLAST alignment and data processing

We firstly build up BLAST databases using all genome assemblies (command: *makeblastdb* − *in genome_assembly_name* − *dbtype nucl* − *out database_name*). Then, BLAST alignments were carried out by querying all annotated IFN-I towards the BLAST databases (command: *tblastn* − *query IFN_input* − *db db_name* − *out alignment_result_name − evalue* 1*e* − *outfmt* 0). Next, all BLAST alignment result sequences were examined forward and backward (under open reading frame) along their genomic locations to identify sites that corresponding to the start codon (i.e., ATG) and the stop codon (i.e., TAA, TAG, and TGA), and corresponding sequences (start codon to stop codon) are saved as test dataset. They are subsequently input into the IFN-SCOPE model to have prediction results for unannotated IFN-I.

#### 1.3 Results filter by IFN-SCOPE model

##### 1.3.1 Data extraction and training dataset construction

We extract all annotated IFN-I **(Materials & Methods 1.1.3)** and their neighborhood-genes from the gff files as the candidate genes for investigation. Here, IFN-SCOPE model is detailed by **Figure S2**. Here, we define the term “neighborhood-genes” as “10 genes (IFN-I genes are excluded) on the same chromosome that are closest to the position of a particular IFN-I gene A (*A_position_*), which is previously reported^15^. *A_position_* is described by **Eq. 1**

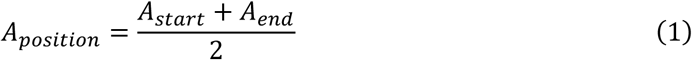

where *A_start_* and *A_end_* denote the start and end site of gene A on the chromosome, respectively. Subsequently, we use the amino acid sequences of all IFN-I and their neighborhood-genes to construct the training dataset. Here, the amino acid sequences of some IFN-I neighborhood-gene cannot be successfully accessed or downloaded due to the non-standardization of the database. They have not been included in our study.

##### 1.3.2 Feature selection

We do feature selection^16–18^ through amphiphilic pseudo amino acid composition (APAAC)^19^ method. Suppose an IFN-I amino acid sequence *P* is composed of *L* residues, which is described by **Eq. 2**

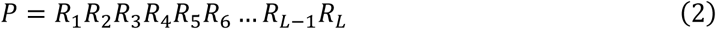

where *R*_*n*_(*n* = 1,2,3, …, *L*) denotes the *n* − *th* amino acid of *P*, *R*_*n*_ = *Alanine*(*A*), *Leucine*(*L*), *Glycine*(*G*) …. Then, we extract the following 20 + 2*λ* dimensional feature vector *V* from *P* (we adopted *λ* = 30 in this study from previous report^20^), described by **Eq. 3**

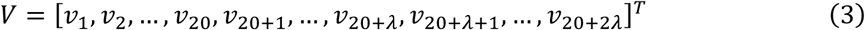

where *T* is the transpose sign, *v*_*u*_(*u* = 1,2,3, …,20 + 2*λ*) is the *u* − *th* feature in *V*, which could be computed by **Eq. 4**

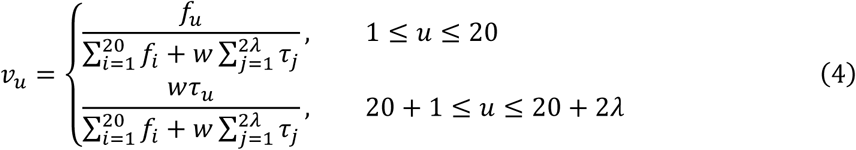

Here, *w* is the weight factor (we adopted *w* = 0.05 in this study from previous report ^20^); *f*_*i*_ (*i* = 1,2,3, …,20) is the normalized occurrence frequency of the *i*-th amino acid in *P*, which reflects the compositional information of IFN-I amino acid sequence; *τ*_*j*_(*j* = 1,2,3, …, 2*λ*) denotes the *j* − *tier* hydrophilicity-hydrophobicity factor of *P*, which reflects the sequence-related

structural information of IFN-I and is computed by **Eq. 5**

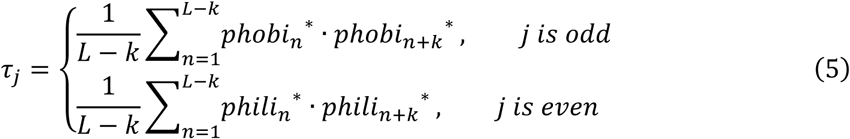

Here, *k* = 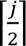 (rounding off of 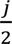); *phobi*_*n*_^∗^ and *phili*_*n*_^∗^ respectively denote the normalized hydrophobicity value and normalized hydrophilicity value of the *n*-th amino acid *R*_*n*_, and these values can be obtained from Z-Score normalization based on previous reported hydrophobicity and hydrophilicity values^21,22^, which are described by **Eq. 6** and **Eq. 7**. The dot (·) denotes the multiplication sign.

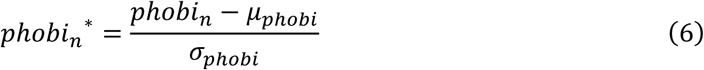

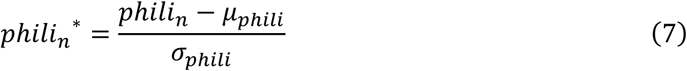

Here, *phobi*_*n*_ and *phili*_*n*_ denote the hydrophobicity and hydrophilicity values of *R*_*n*_, respectively; *µ*_*phobi*_ and *µ*_*phili*_ denote the mean values of all *phobi*_*n*_ and *phili*_*n*_ values, respectively; *σ*_*phobi*_ and *σ*_*phili*_ denote the standard deviation of all *phobi*_*n*_ and *phili*_*n*_ values, respectively.

##### 1.3.3 Feature dimensional reduction

We did feature dimensional reduction^23–25^ based on the feature importance of the model^26^, which aims at to maximize the predictive accuracy of IFN-SCOPE model. Specifically, the model is firstly trained by initial set of features, and then we obtained the coefficient *ω*_*u*_ for each feature *x*_*u*_. After that, feature *x*_*u*_ would be considered unimportant and purned if the absolute value of their corresponding coefficient |*ω*_*u*_| was less than the threshold 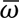 (**Eq. 8**). In this study, we carried out SelectFromModel algorithm^26^ for dimensional reduction.

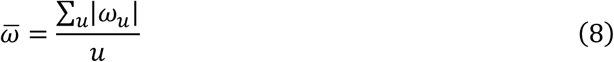

Here, 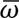 is the average of all |*ω*_*u*_|, *u*(*u* = 1,2,3, …,80) is the index of the feature *x*_*u*_. A k-fold cross-validation is carried out on the dataset before and after dimensional reduction, respectively, and the average of *F*_1_-score **(Eq. 9)** is used to evaluate the performance of the feature dimensional reduction.

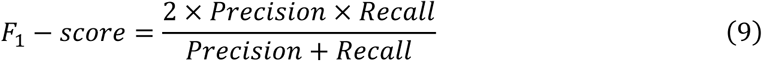

Here, *Precision* and *Recall* can be obtained from **Eq. 10** and **Eq. 11**, respectively.

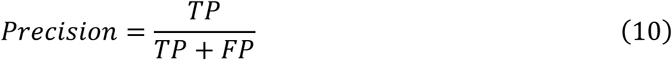

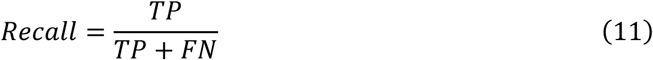

Here, *TP* (true positive) is the number of perfectly identified IFN-I, *TN* (true negative) is the number of perfectly identified IFN-I neighborhood-genes, *FP* (false positive) is the number of wrongly identified IFN-I which is actually IFN-I neighborhood-genes, *FN* (false negative) is the number of wrongly identified IFN-I neighborhood-genes which is actually IFN-I.

##### 1.3.4 Model construction and training

We used the logistic regression^27–30^ as the classifier of our model **(Figure S2)**. By adopting the L2-regularization, logistic regression reduces the complexity of coefficients, and thus prevent model from overfitting. Specifically, we used feature vectors as the input, which is the output of the feature dimensional reduction procedure, and used the sample labels (is/not IFN-I) as the output. Our logistic regression was optimized by a stochastic average gradient descent algorithm.

##### 1.3.5 Model evaluation and prediction

We used *F*_1_-score (**Eq. 9**), *Accuracy* (**Eq. 12**) and area under *ROC curve* (**Eq. 13**) to evaluate the performance of the IFN-SCOPE model.

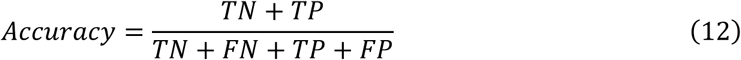

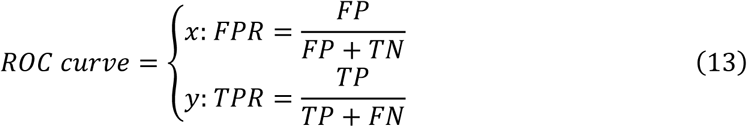

In **Eq. 13**, *FPR* and *TPR* denote the false positive rate and true positive rate of the model under different thresholds, respectively. Subsequently, the well-trained IFN-SCOPE model is used to predict the label for each sample in the test dataset **(Materials & Methods 1.3.1)**. Samples with positive predictions are saved as unannotated IFN-I after double-filtering and reviewed by annotated IFN-I, and they are finally named as ‘ *species_chromosome_position* ’ (such as *Anas_platyrhynchos_NC_051804.1_POS71719872*) **(Table S1)**.

### 2. IFN-I conserved loci computation and assignment

#### 2.1 Gene conserved loci computation

The degree centrality algorithm^31,32^ indicates the node with the maximum degree (highest number of direct connections) is most important. Here, we developed the GENE-GRADE algorithm to look for the conserved loci within IFN-I evolution of all genes for our candidate species, which consists of two parts: knowledge graph construction and conserved loci computation. Here, we detail the GENE-GRADE algorithm by **Figure S3**.

##### 2.1.1 Knowledge graph construction

We only considered genes that are in the revised GFF files and placed on the chromosome of our candidate species during knowledge graph^33–35^ construction **(Materials & Methods 1.1)**. Then genes are added into knowledge graph *G* as individual nodes *N*_*gene*_, and merged IFN-I nodes which are adjacent to the gene position **(Materials & Methods 1.1.2)** in *G*. Next, we build up the gene chains by mergeing IFN-I nodes *N*_*IFN*_ and IFN-I neighborhood-genes *N*_*nb*_*gene*_ according to their gene position. Finally, we remove all nodes that were not placed on gene chains.

##### 2.1.2 Conserved loci computation

We merge all *N*_*nb*_*gene*_ which have the same gene name for all gene chains in *G*. Then, the degree centrality algorithm is employed to look for the node with the maximum degree. If the degree of *N*_max _*degree*_ is greater than 15, the gene of *N*_max _*degree*_ will be outputted and saved as a conserved locus, after that we remove all gene chains that include *N*_max _*degree*_ in *G*; else, we conside that the gene of *N*_max _*degree*_ is existed only in some of our candidate species, which contradicted the meaning of the term “conserved”, and the GENE-GRADE algorithm ends.

#### 2.2 Conserved loci assignment

After computing the conserved loci of IFN-I evolution by GENE-GRADE algorithm, we assign IFN-I genes into their corresponding conserved locus according to the definition of neighborhood-genes **(Materials & Methods 1.3.1)**. If one IFN-I gene could not be assign into any conserved locus, we assign it into the “exception” category.

The visualization of all conserved loci, gene orientations, numbers, and neighborhood-genes results of IFN-I genes are implemented by ggplot2 and gggenes packages in R, as well as seaborn in Python. Specifically, we adopt following principles in presenting our results. First, if the nomenclature of a IFN-I in the public database is consistent with our analysis results, the name will be displayed above its gene arrow; else, the name that displayed inside the parentheses represents the nomenclature from our results, while the name that displayed outside the parentheses represents the nomenclature in public database **(Figure S7 & S8)**. Second, the relative position of gene arrows at the same conserved locus are computed based on the position **(Materials & Methods 1.3.1)** of IFN-I genes.

### 3. IFN-I comparative genomics evolutionary analysis

#### 3.1 Multi-sequence alignment

We carry out multi-sequence alignments (MSA)^36^ in this study after inputting the .fasta files of interest into MAFFT (v7.520) (command line: *mafft* − −*auto* − −*reorder* − −*leavegappyregion input*_*filename* > *output*_*filename*). The visualization of results are implemented by ggplot2, and ggmsa packages in R. In sequence identity matrices, species categories are colored that corresponds to the taxon in the phylogenetic tree. The ends of the MSA result may be trimmed for clearer presentation.

#### 3.2 Phylogenetic analysis

Phylogenetic trees^37^of IFN-I are constructed through the MSA **(Materials & Methods 3.1)** results of interest by MEGA. All trees of IFN-I are tested by bootstrapping (1500 trials), with the results indicated by the color of the clade nodes (red for *bootstrap value* > 90, pink for 90 > *bootstrap value* > 70, and white for *bootstrap value* < 70). We adopt Ig domain containing 2 (LINGO2) proteins from candidate species as outgroups of IFN-I trees in this study. Phylogenetic trees of species are constructed through TimeTree^38^.

The visualization of all phylogenetic results is implemented by ggplot2, ggtree, and ggstar packages in R. Specifically, we adopted the following principles for assigning or giving nomenclature to unnamed IFN-I in trees. First, for previously reported IFN-I that from placentals, we name all IFN-Is within the same monophyletic group based on previous name of their subtype (such as α, ω, β); for previously unreported IFN-I that mainly presented in stem mammals, birds, and reptiles, we propose new nomenclatures (such as H2, H1, U2, and U1) based on the evidence from synteny, sequence homology, and copy number. Low quality of IFN-I node that is not group with any other lineages will be purned for clarity **(Table S3)**.

#### 3.3 Sequence identity computation

We computed identity values between all amino acid sequence of IFN-Is through ClustalO. All amino acid sequences are firstly input (command line: − − *infile input*_*filename* − *threads* 8 − −*MAC* − *RAM* 8000 − −*verbose* − −*full* − −*outfmt clustal* − −*resno* − −*outfile output*_*filename* − −*output* − *order tree* − *order* − −*seqtype protein*) and then converted to .csv format for accessing identity values between amino acid sequences. The visualization of identity matrix results is implemented using the Seaborn in Python.

### 4. IFN-I functional experiment

#### 4.1 PCR amplification, plasmid construction, and expression of IFN-Is

We extract DNA from the anticoagulated blood of baer’s pochard (*Aythya baeri*) by TIANamp Genomic DNA kit (DP304, TIANGEN). Next, we designed primers **(Table S2)** to carry out PCR amplification and sequencing analysis for baer’s pochard IFN-CL gene and its flanking sequences, which includes a 22,432 bp DNA segment that from the COG6 to LHFPL6. Then, gene fragments of IFN-CL and IFN-U2 are cloned into PTT3 vector with a C-termianl His tag. All the plasmids are transfected into HEK293F cells and cultured in a constant temperature shaker with 8% CO2 at 37 °C. After 96 hours of culture, the supernatant is then collected by centrifugation for 10 minutes and used for quantitative Western Blot analysis.

#### 4.2 PBMC isolation and IFN-I stimulation

Baer’s pochard PBMCs are prepared from anticoagulated blood by duck peripheral blood lymphocyte isolation kit (LTS1090D, TBD Sciences). The cells are maintained in RPMI 1640 medium (Thermo Scientific) with 10% FBS at 37°C with 5% CO2. Next, PBMCs are stimulated with IFN-CL and IFN-U2 for 6h.

#### 4.3 Quantification of ISGs by RT-PCR

Stimulated cells are collected and washed in ice-cold PBS, and then pelleted and lysed with 0.5 ml TRIzol (15596026CN, Invitrogen). Total RNA from cells is extracted and then treated with RQ1 RNase-Free DNase (M6101, Promega), and reverse-transcribed in 25 μl reactions using M-MLV Reverse Transcriptase (M1701, Promega) according to the manufacturer’s instructions. To quantify transcript levels, 20 μl reactions are set up containing 1 μl reverse transcription product, 2× PowerUp SYBR Green Master Mix (A25742, Applied Biosystems), 250 nM forward and reverse primers **(Table S2)** of glyceraldehyde 3-phosphate dehydrogenase (GAPDH) or 2’-5’-oligoadenylate synthetase like (OASL). The following cycling conditions on the QuantStudio Q6 (Applied Biosystems) system are used: 10 min at 95 °C; then 40 cycles of 15 s at 95 °C and 1 min at 60 °C.

## Results

### 1. 115 unannotated IFN-Is are identified by IFN-SCOPE model

The frequent duplication and rapid diversification leads to low sequence homology of IFN-I genes between evolutionaily distant animals, or even between different IFN-I subtypes in the same animal species. The low sequence homology of IFN-Is hampers the complete and accurate annotation of IFN-I genes. Therefore, we develop a computational method based on BLAST^14^ and a custom IFN-SCOPE model **(Materials & Methods 1)** to systematically identify unannotated type I interferon (IFN-I) genes in candidate species.

First, we extract 793 annotated IFN-I genes from 87 amniotes **(Figure S1) (Materials & Methods 1.1)**. These IFN-Is include 56 IFN-α and 15 IFN-β (in mammals), 24 IFN-κ (in mammals and birds), 12 IFN-ε (in mammals), 11 IFN-δ (in sheep), 10 IFN-ω (in human, artiodactyls and chicken), 7 IFN-τ (in ruminants), 1 IFN-ζ (in mouse), 14 IFN-Is that named with “Gm” (such as Gm13282, in mouse), 5 IFN-Is that named with numbers (such as IFN1, in platypus and sheep), and 638 unclassified IFN-I genes that identified with “LOC” numbers. All these IFN-I genes have amino acid sequences provided by public databases^9^.

Next, we use amino acid sequences of annotated IFN-Is as query sequences, and obtain potential protein-coding regions of IFN-I on the genomes of candidate species by BLAST **(Materials & Methods 1.2)**. Then, we input newly identified sequences into the IFN-SCOPE model to have predictive results for unannotated IFN-I **(Materials & Methods 1.3)**. A total of 793 positive samples (i.e., amino acid sequences of IFN) and 2041 IFN-I adjacent genes are extracted into the training dataset after data preprocess.

Next, we carry out feature selection based on the amino acid composition, the hydrophobicity and the hydrophilicity correlation between residues of sample sequences **(Materials & Methods 1.3)**. After that, we did feature dimensional reduction to decrease the overfitting risk and improve the model performance. As presented, the feature space is reduced from 80 dimensions to 36 dimensions after dimensional reduction (**Table 1**). Subsequently, matrices and corresponding sample labels are input into a logistic regression classifier for model training, and we evaluate the performance of the model before and after the feature dimensional reduction by various metrics of a 5-fold cross-validation (**Table 1**). All of the model metrics improved after feature dimensional reduction, which demonstrates the effectiveness of the procedure.

**Table 1.** Feature dimension, F_1_-score, accuracy, and area under ROC curve(AUC) before and after feature dimensional reduction.

| Metrics | Before dimensional reduction | After dimensional reduction |
| --- | --- | --- |
| Feature numbers | 80 | 36 |
| F<sub>1</sub>-score | 0.9816 | 0.9830 |
| Accuracy | 0.9901 | 0.9908 |
| AUC | 0.9934 | 0.9945 |

Finally, we input the test set that constructed by BLAST into our well-trained logistic regression model to obtain the predictive results for unannotated IFN-Is. 115 unannotated IFN-Is are discovered by the IFN-SCOPE model **(Table S1)**.

### 2. Three conserved IFN-I loci computed by GENE-GRADE are located on the same chromosome

After obtain all IFN-I genes, we develop a GENE-GRADE algorithm **(Materials & Methods 2.1)** for computing the conserved loci of amniote IFN-I genes based on the degree centrality algorithm^31^. Due to the short-length and incomplete gene annotation of unplaced genome scaffolds, we only consider IFN-I genes placed onto chromosomes of the candidate species for conserved loci analysis, which includes 556 annotated and 75 unannotated IFN-I genes.

The aforementioned 631 IFN-I genes are processed with GENE-GRADE algorithm, and subsequently output three (HACD4, MOB3B and UBAP2) conserved loci, which are named after the adjacent genes of the IFN-I clusters (**Figure 2A**). IFN-I genes are located at those three conserved loci constitute 95.7% (604 out of 631) of all IFN-I genes in our study (**Figure 2B**), and the majority of them localize to the HACD4 locus (445 annotated, 35 unannotated), while fewer reside at UBAP2 (71 annotated, 15 unannotated) and MOB3B (19 annotated, 19 unannotated) **(Materials & Methods 2.2)**. The remaining 27 genes (21 annotated, 6 unannotated) assign to uncharacterized loci (‘exception’ category) or require further validation.

**Figure 2.**
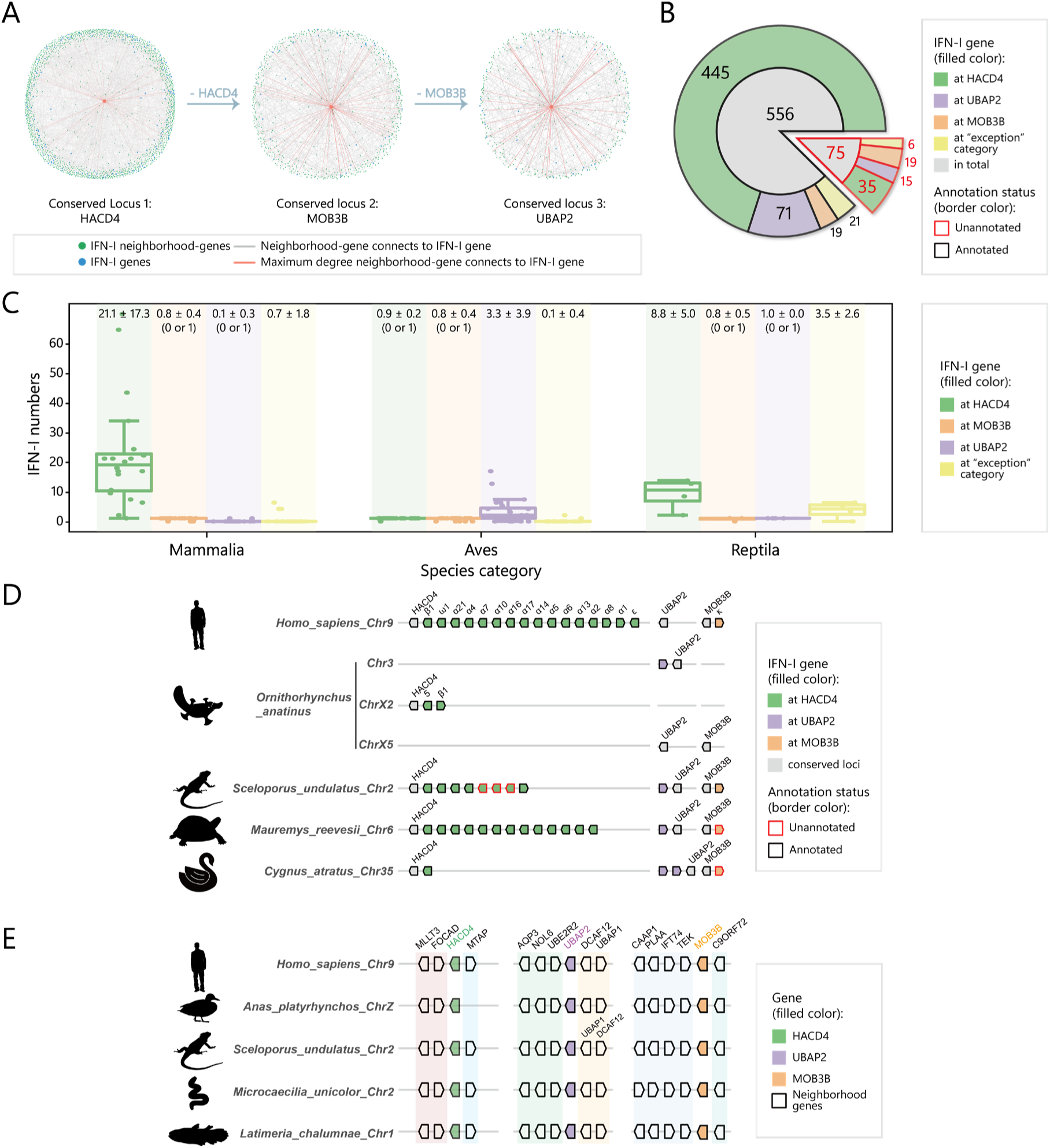
The Conserved loci computed from GENE-GRADE algorithm. **(A)** The result of the GENE-GRADE algorithm. Each subplot represents the knowledge graph before outputting conserved loci HACD4, MOB3B, and UBAP2, respectively. Gray lines represent the connections between nodes in the knowledge graph, whereas the orange lines represent the connections between maximum degree node (the node of conserved locus) and other nodes. **(B)** The numbers of placed IFN-I gene on conserved loci or “exception” category. IFN-I genes are assigned into annotated or unannotated in the pie chart, and their corresponding numbers are shown in the inner circle of the pie chart; while the outer circle illustrates the numbers of IFN-I gene at HACD4 locus, UBAP2 locus, MOB3B locus, and “exception” category. **(C)** Box plots of the numbers of IFN-I genes at conserved loci or “exception” category for species in mammalia, aves, and reptila. The label on the top of diagrams represent the mean and standard deviation of the corresponding boxes, and ‘(0 or 1)’ will be marked behind the statistical value if there is only a single-copy IFN-I gene or no IFN-I gene at corresponding locus. **(D)** The distribution of all IFN-I genes at conserved loci or “exception” category in representative species. IFN-I genes on the chromosomes of human *(Homo sapiens)*, platypus *(Ornithorhynchus anatinus)*, fence lizard (*Sceloporus undulatus*), turtle *(Mauremys reevesii)*, and black swan (*Cygnus atratus*) are displayed according to their corresponding conserved loci. **(E)** The distribution of three conserved loci and their neighborhood-genes in four species. Conserved neighborhood-genes are marked by different color shadings. IFN-I genes are not shown, and the gene orientations are adjusted to align the gene orientations in human in this Figure.

Interestingly, the numbers of IFN-I genes at 3 conserved loci varies among different animal groups (**Figure 2C, Figure S4, Figure S5, & Table 2**). In all the animals analyzed, IFN-I genes are monogenic or absent at MOB3B locus (**Figure 2C, 2D, & Table 2**). Even if IFN-I genes at HACD4 locus are monogenic in birds, but they are polygenic in mammals (median: 21) and in reptiles such as turtles, crocodilians, and squamates (median: 9) (**Figure 2C & 2D)**. Conversely, IFN-I genes at UBAP2 locus are monogenic in mammals and reptiles but polygenic in birds (median: 3) (**Figure 2C & 2D)**, but in mammals, only monotremes (platypus and echidna) have a single-copy IFN-I gene at UBAP2 locus (**Figure 2D & Figure S4)**.

**Table 2.**
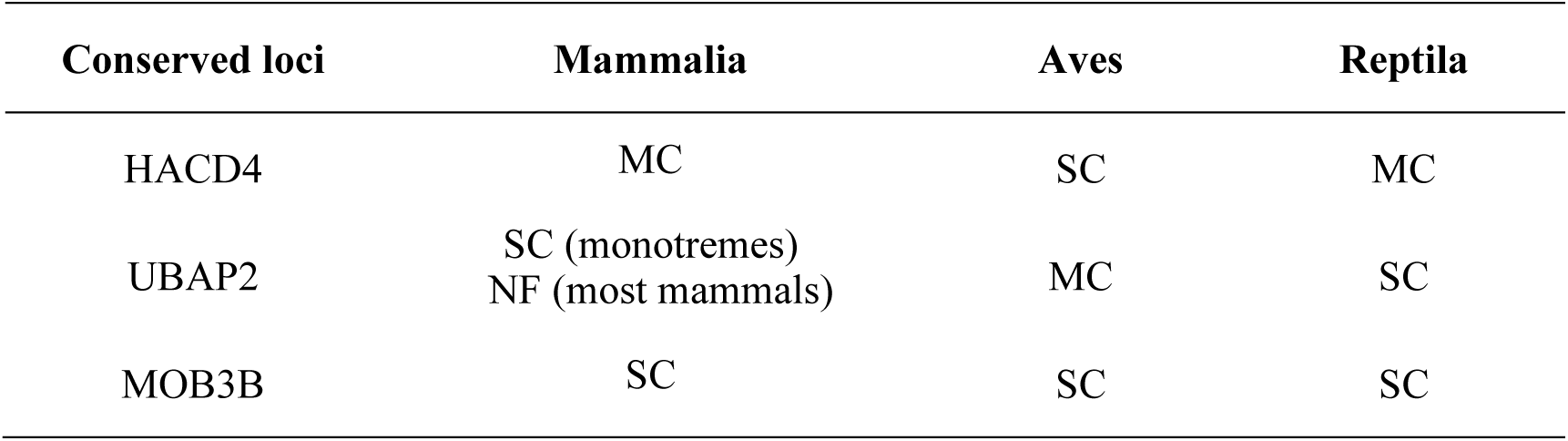
The numbers of IFN-I genes in the conserved loci. MC: multiple-copy; SC: single-copy; NF: not found.

The DNA fragments containing those 3 conserved IFN-I loci (HACD4, MOB3B and UBAP2) locate on the same chromosome in most animals we analyzed, with few exceptions including pig (*Sus Scrofa*), common brushtail possum (*Trichosurus vulpecula*), platypus (*Ornithorhynchus anatinus*) and echidna (*Tachyglossus aculeatus*) (**Figure 2D, Figure S4, & Figure S5)**. Importantly, the conserved colocalization of HACD4, MOB3B, and UBAP2 on a single chromosome predates the integration of intronless IFN-I genes, as evidenced in unstriped caecilian (*Microcaecilia unicolor*) and coelacanth (*Latimeria chalumnae*) (**Figure 2E**). Collectively, these observations support the hypothesis that intronless IFN-I genes were already integrated at the HACD4, MOB3B, and UBAP2 loci in the most recent common ancestor (MRCA) of modern amniotes.

### 3. Both evolutionary trajectories and locus-specificity contribute to the complexity of IFN-I phylogeny

We perform phylogenetic analysis of collected amniote IFN-I genes **(Materials & Methods 3.1 & 3.2)**, and plotted the data with both species and locus information. Surprisingly, we find that the phylogenetic tree displayed a mosaic pattern, reflecting a mixture of evolutionary history and locus-specificity (**Figure 3A & 3B)**.

**Figure 3.**
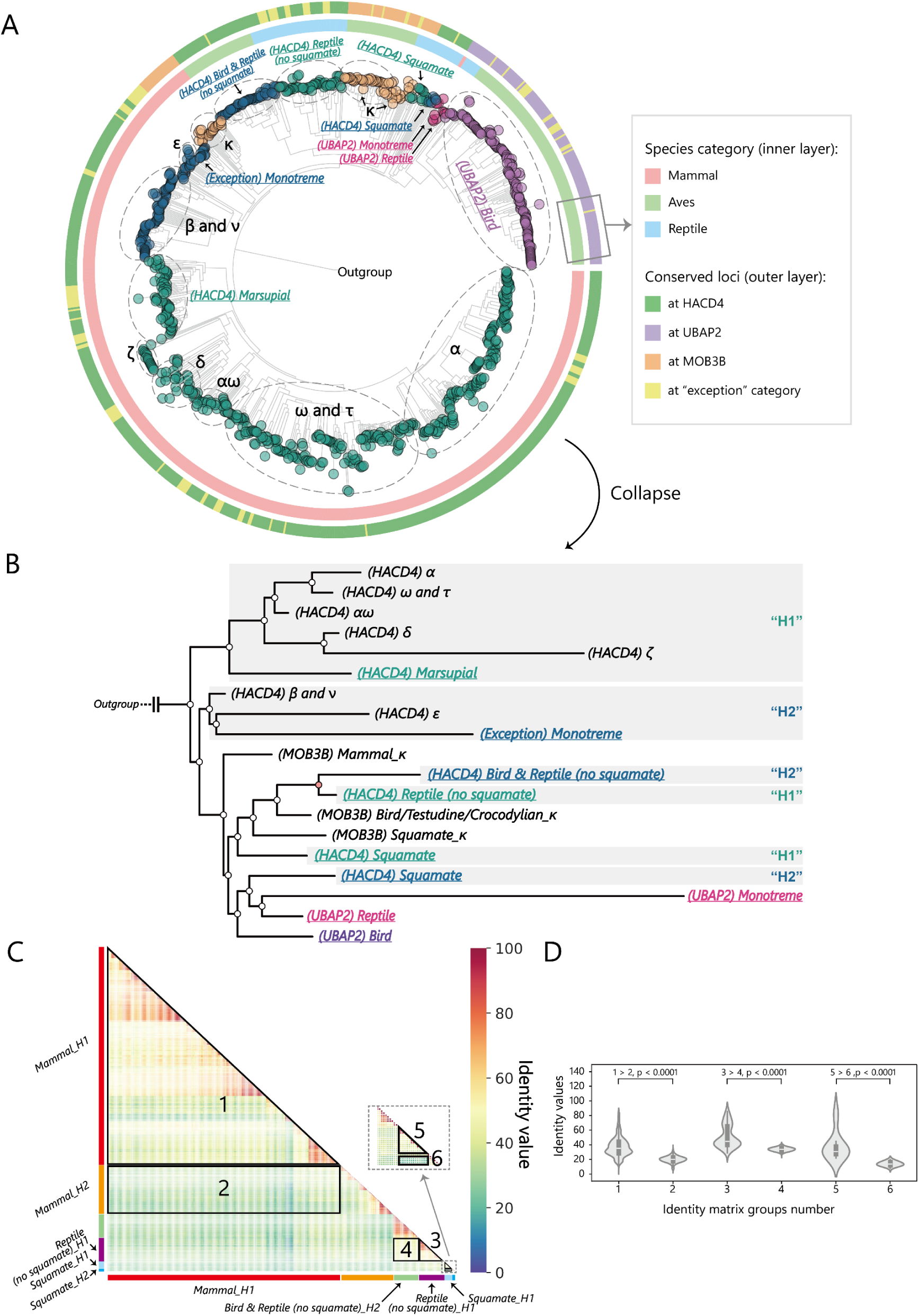
The phylogenetic relationships of amniote IFN-I genes. **(A)** Amniotes IFN-I genes can be assigned into the following clades: H1(dark green), H2 (dark blue), κ (brown), U1 (dark purple), and U2 (light purple). Layers that surrounding the tree represents the corresponding species category and the conserved loci, respectively. First-reported IFN-I clades are highlighted by underlining and named by conserved loci and species category in the figure. **(B)** The phylogenetic tree that collapsed according to the IFN-I clades of in **(A)**. The order of taxa in the phylogenetic tree, from top to bottom, corresponds to the clockwise order of tree taxa in **(A)**. **(C)** The sequence identity matrix in phylogenetic tree of **(A)** and **(B)**. The grouping of taxa is indicated by color bars. **(D)** Violin plots of identity values in the six regions in **(C)** and the significance test results.

Specifically, all IFN-I genes at UBAP2 locus (IFN-U) are almost clustered together (**Figure 3A & 3B)**. The IFN-I genes at MOB3B locus are phylogenetically classified into three distinct lineages: mammals, squamates, and Archelosauria (including turtles, crocodilians, and birds) (**Figure 3B**). A parallel evolutionary pattern is observed at HACD4 locus, where these IFN-I genes (IFN-H) similarly diverge into these three lineages.

Notably, each lineage-specific IFN-H group further diversifies into two subclusters: the first cluster of mammalian IFN-Hs (mammalian IFN-H1, **Figure 3B**) comprises all multi-copy subtypes (IFN-α, IFN-ω, IFN-τ, IFN-αω, IFN-δ, IFN-ζ) of placental mammals^3,9,10,39^, as well as multiple-copy IFN-I genes of marsupials. The second cluster of mammalian IFN-Hs (mammalian IFN-H2, **Figure 3B**) includes IFN-β, IFN-υ, and IFN-ε of therian mammals, along with all IFN-I genes of monotremes^9^. Correspondingly, the Archelosaurian IFN-H1 and IFN-H2 clusters are located adjacent to each other in the phylogenetic tree. In contrast, the squamate IFN-H2 genes cluster with IFN-I genes at UBAP2 locus, while the squamate IFN-H1 cluster groups with other IFN-Hs (**Figure 3A & 3B)**.

The division of IFN-H1 and IFN-H2 groups is further supported by sequence identity matrix analysis **(Materials & Methods 3.3)**. The identity values within IFN-H1s are greater than those between IFN-H1s and IFN-H2s in mammals (1 *vs.* 2), Archelosaurians (3 *vs.* 4), and squamates (5 *vs.* 6) (**Figure 3C & 3D)**.

### 4. Evolutionarily conserved IFNK orthologs across amnioates

The MOB3B locus shows remarkable conservation, maintaining one IFN-I copy (or zero) throughout amniote evolution (**Figure 2D, Figure S4, & Figure S5)**, and they form one single cluster among mammialian IFN-I genes (**Figure 4A**). In addition, the neighborhood-genes, such as C9ORF72, LINGO2, and TEK, are conserved in most amniotes **(Figure S6A)**. Notabely, all IFN-I genes at MOB3B locus are co-oriented and transcribed in the same direction with MOB3B **(Figure S6A)**. Based on synteny analysis, we propose that the IFN-I genes at MOB3B locus represent orthologs of human IFNK (coding gene of IFN-κ) throughout amniotes **(Figure S7 & Figure S8)**. It’s worth pointing out that the IFN-I gene at HACD4 locus of chicken (*Gallus gallus*) and mallard (*Anas platyrhynchos*) have been classified as the IFN-κ based on amino acid sequence homology^40,41^. However, both of animals have a truncated form of IFNK sequence beside MOB3B, argue against the nomenclature of IFN-I at HACD4 locus of birds as the orthologs of human IFNK **(Figure S8)**.

**Figure 4.**
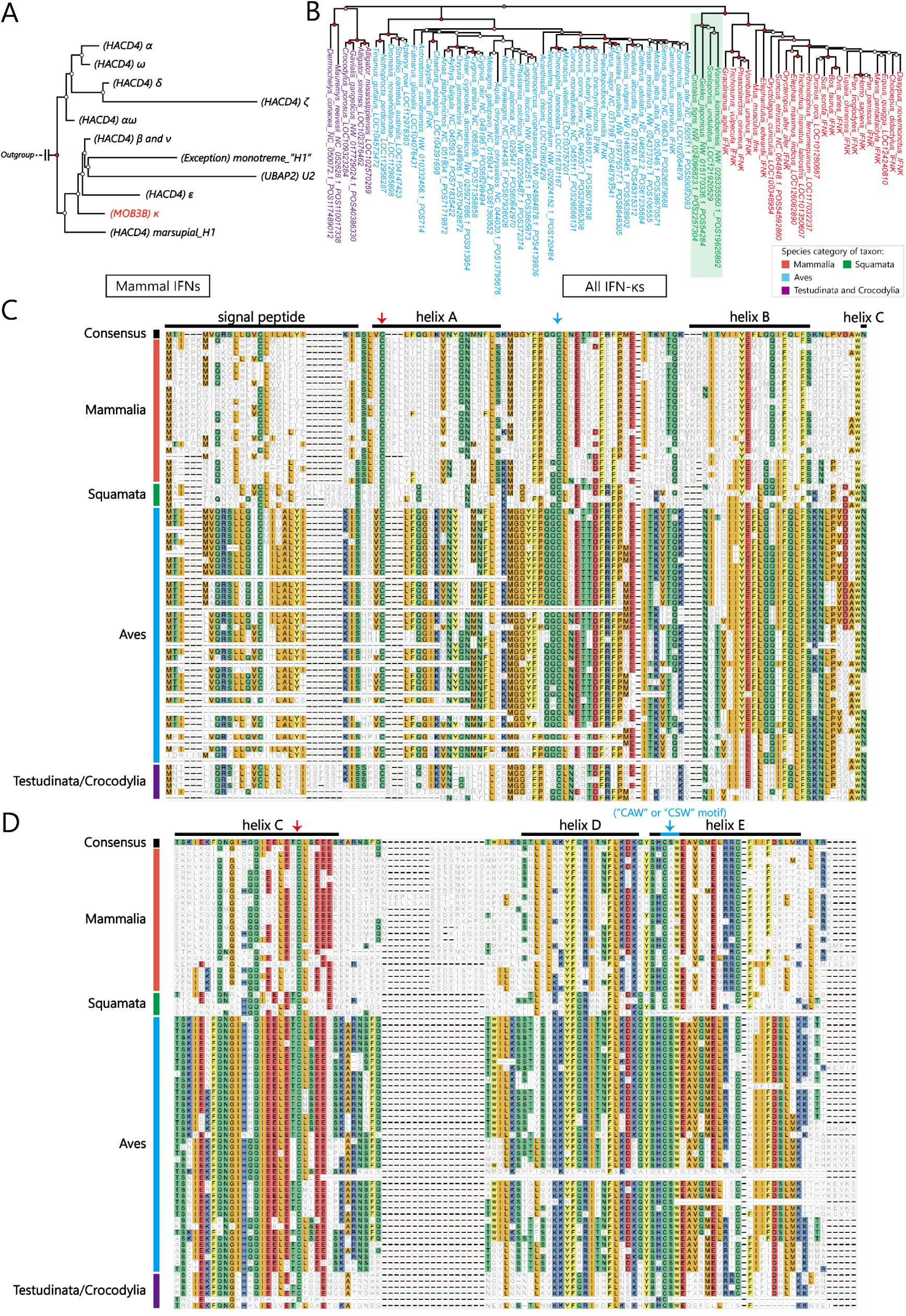
The evolution of IFN-κ orthologs. **(A)** The phylogenetic relationships of all mammal IFNs. The monophyletic group of IFN-κs is colored in red. **(B)** The phylogenetic relationships of all IFN-κs. Species from mammalia (red), squamata (green), aves (blue), and testudinata & crocodylia (purple) are respectively colored. **(C)** The first half MSA result of all IFN-κs in our candidate species. **(D)** The second half MSA result of all IFN-κs in our candidate species.

Interestingly, our results show that IFN-κ of squamates (green colored in **Figure 4B**) form a monophyletic group with mammals’ IFN-κ first, and then they groups with the remaining birds and reptiles IFN-κ. Moverover, the IFN-κ of birds, testudines, and crocodylians is greatly different from those in mammals and squamates, as indicated by the filled-color of the MSA results (**Fig. 4C**). Additionally, around the fourth conserved cysteine^10^ (blue arrow in **Fig. 4D**), mammalian and squamate IFN-κ share a conserved ‘CAW’ motif (cysteine-alanine-tryptophan), while avian, testudine, and crocodylian IFN-κ exhibit a ‘CSW’ motif (cysteine-serine-tryptophan). Thus, we propose that the major speciation event of IFN-κ might occur during the squamata-testudinata speciation, rather than the mammalia-sauropsida speciation as indicated by taxonomy **(Figure S1)**.

### 5. Two-subcluster division of IFN-Hs are conserved between mammals and reptitles

Both mammals and reptiles possess multiple IFN-Hs while birds contain a single IFN-H gene at HACD4 locus (**Figure 2 & Table 2**). Phylogenetic analysis of all IFN-H genes demonstrates that squamate IFN-Hs cluster together and form a sister group to mammalian IFN-H2s (IFN-β/υ/ε), with mammalian IFN-H1s (IFN-α/ω/δ) diverging earlier (**Figure 5A**). Notably, the multiple-copy monotreme IFN-H1 genes (**Figure 5A**, star-labeled) are located on Chromosome X5 **(Figure S7)**, where they neighbor TBC1D2 gene that typically flank MOB3B in other vertebrates **(Figure S6)**. However, unlike canonical IFNKs, they are not positioned between MOB3B and C9ORF72 **(Figure S7)**. Phylogenetically, these genes cluster with IFN-Hs rather than IFNKs, suggesting their evolutionary origin as true IFN-H homologs. This unique genomic arrangement of monotreme IFN-H1s likely reflects specialized ecological and microbial adaptations of these egg-laying mammals^42^.

**Figure 5.**
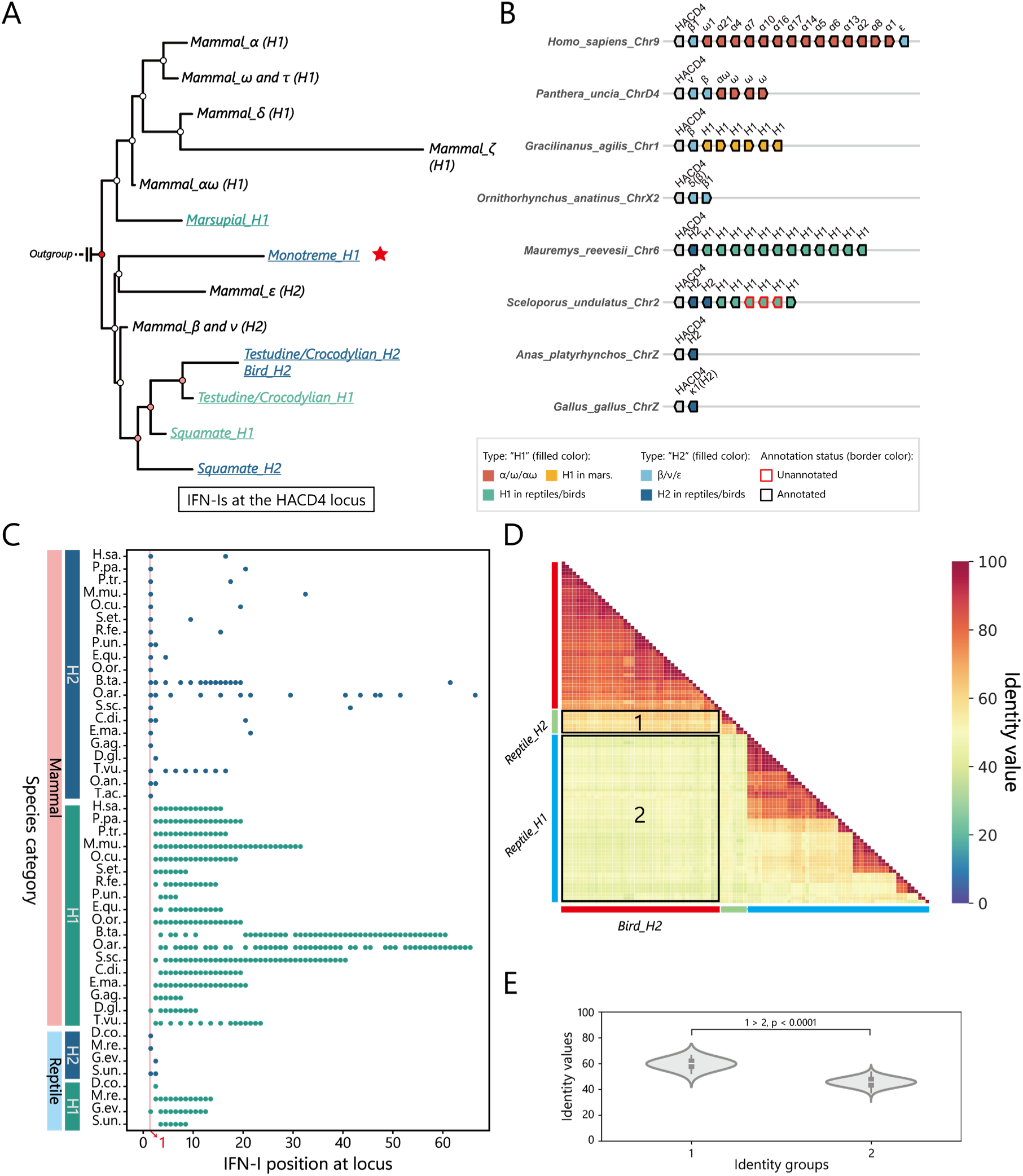
IFN-Hs can be classified as H1 and H2. **(A)** The phylogenetic relationships of all IFN-Hs. H1 or H2 are marked for every corresponding clade in the tree. First-reported IFN-I clades are highlighted by color and underlining. A red star indicates the specific IFN-H1 clade in monotremes. **(B)** The distribution of IFN-I genes at HACD4 on the chromosomes of representative species. IFN-I subtypes can be classified into “H1” and “H2”. **(C)** The relative IFN-I position of IFN-Hs. Each data point represents the position of an IFN-I gene at HACD4 in the corresponding species. **(D)** The sequence identity matrix of H1 and H2 genes in reptiles and birds. The grouping of taxa is indicated by color bars. **(E)** Violin plots of identity values in the two regions in **(D)** and the significance test results.

In mammals and reptiles, the IFN-H genes diverged into IFN-H1 and IFN-H2 subclusters (**Figure 3 & Figure 5A**). Notably, HACD4-adjacent IFN-H genes are phylogenetically restricted to the IFN-H2 subcluster and in the same transcriptional orientation (**Figure 5B & 5C)**. Further analyses reveal a consistent pattern across species: the HACD4-adjacent IFN-H2 gene is typically maintained as a single-copy (e.g., IFN-β), while IFN-H1 genes frequently undergo multiple-copy expansion (**Figure 5B & 5C)**.

All birds possess only one IFN-I gene at HACD4 locus. Sequence identity matrix reveals this avian IFN-H shares statistically significant greater identity with mammalian IFN-H2s than with IFN-H1s (p< 0.0001; **Figure 5D & 5E**). These data suggested that the single-copy IFN-H2 gene represents the ancestral state at HACD4, and mammalian IFN-H1 clusters evolved via lineage-specific duplications in mammals and reptiles.

### 6. Convergent diversification of IFN-I genes at UBAP2 locus of birds

Although chicken IFN-Is at UBAP2 locus (IFN-Us) were initially proposed as homologs of human IFN-α/β^43^, syntenic and phylogenetic analyses reveal these genes evolved independently from HACD4-linked IFN-Is^9,44^. Unlike the dichotomous IFN-H1/H2 clusters at HACD4 locus, IFN-Us almost form a single monophyletic group (**Figure 2A & 2B)**, underscoring the distinct evolutionary histroy of IFN-Us comparing with IFN-Hs.

When phylogenetically analyzed as a separate group, IFN-U genes consistently bifurcate into two distinct subgroups (IFN-U1 and IFN-U2) within each avian order we examined (**Figure 6A**). Thus, IFN-I genes at UBAP2 locus have been revised to 44 IFN-U2 and 95 IFN-U1 **(Figure S7 & Figure S8)**. Sequence identity matrix reveals that higher identity values among IFN-U1s within the same order (regions 1, 3, 5) compared to IFN-U1 *vs.* IFN-U2 (regions 2, 4, 6) and this pattern is consistent across different avian orders (e.g., anseriformes, galliformes, and passeriformes) (p<0.0001; **Figure 6B & 6C**). Monotremes and reptiles have a single-copy IFN-U gene, and sequence identity matrix reveals this monotreme/reptile IFN-U shares statistically significant greater identity with bird IFN-U2s than with IFN-U1s (p< 0.01; **Figure 6D**), which proves that they belong to IFN-U2 (**Figure 6A**). Addtionally, the IFN-U2 genes mostly exist in single-copied form and located at the beginning of the IFN-U cluster, whereas the IFN-U1 genes are present in multiple-copied form follows IFN-U2 (**Figure 6E**).

**Figure 6.**
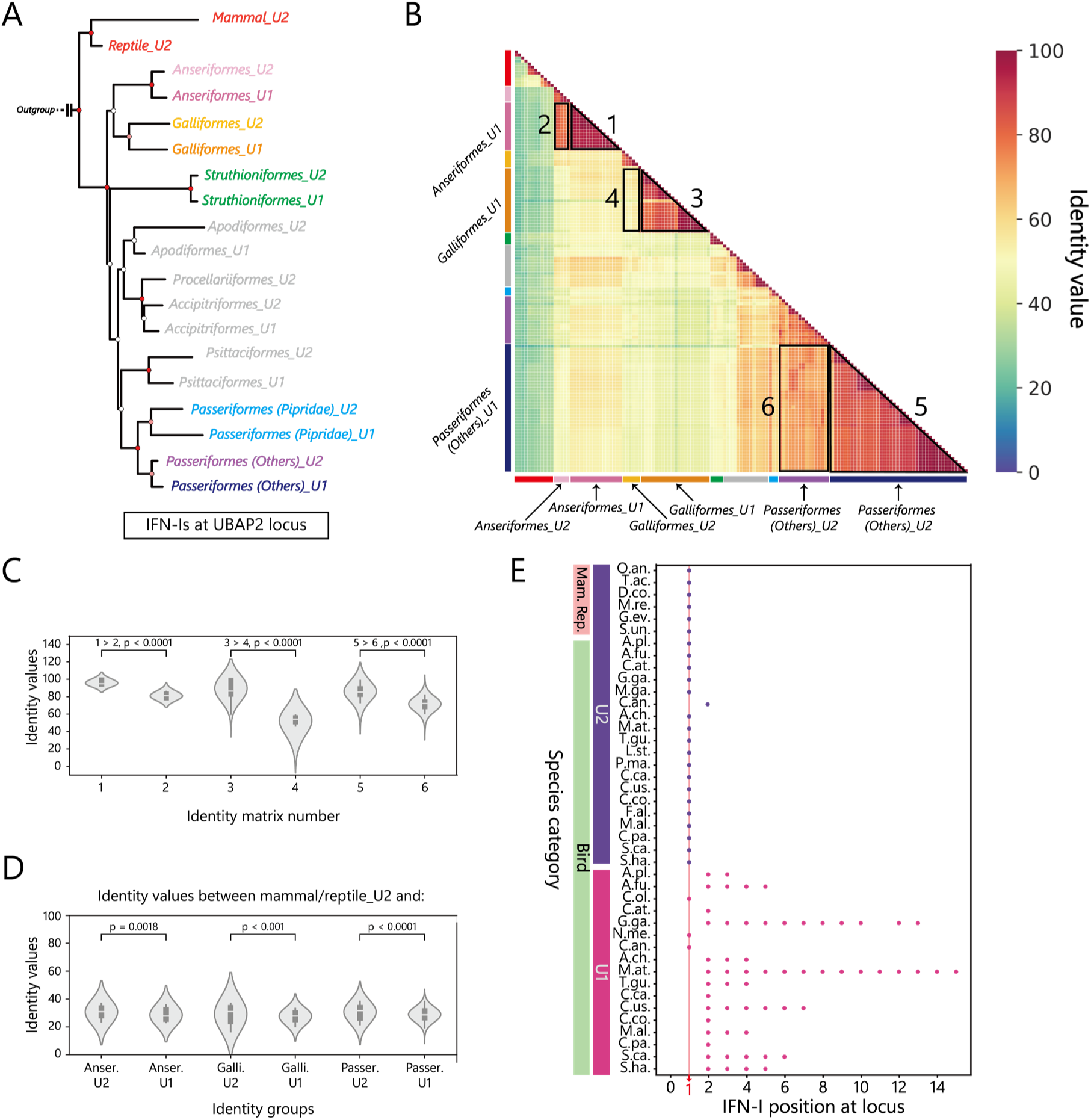
IFN-Us diversify into U1 and U2. **(A)** Phylogenetic relationships of all IFN-Us. Some IFN taxa are colored gray due to the low-quality of the genome assembly of corresponding species. **(B)** The sequence identity matrix of phylogenetic tree in **(A)**. The meaning of each color bar corresponds to the taxon of the same color in **(A)** (i.e. the red bar represents the IFN-U2 in mammals and reptiles). **(C)** Violin plots of identity values in the six regions in **(B)** and the significance test results. **(D)** Violin plots of identity values between mammal/reptile IFN-U2s and other IFN-Us in birds and the significance test results. **(E)** The relative IFN-I position of IFN-Us. Each data point represents the position of an IFN-I gene in the corresponding species.

Although phylogenetically distinct, the IFN-H and IFN-U follow analogous evolutionary trajectories. IFN-Is at both loci originated from single-copy ancestors (IFN-H2 at HACD4; IFN-U2 at UBAP2), and independently diversified into multicopy subtypes via locus-specific duplications.

### 7. Four-cysteines IFN-I is the ancestor of amnionate interferons

While Krause et al. propose IFN-β as the ancestor of mammalian IFN-Is based on phylogenetic analysis^9^, Secombes et al. (2017) argue that 4-cysteine (4C) IFN-Is (e.g., IFN-κ, IFN-U2) represent the ancestral form, given their conservation in fish, amphibians, reptiles, birds, and mammals^10^. However, mammlian IFN-βs only have 2 coserved cysteines (2C), which is unique among all amnionate IFN-Is^10^. The apparent conflict between these competing hypotheses remains unresolved.

In IFN-H2, different form IFN-β, IFN-υ have 4 conserved cysteines, forming 2 disulfy bonds (**Figure 7A**). In contrast to the canonical IFNB-HACD4 arrangement in most therians, we find functional IFNN (encoding IFN-υ) positioned between IFNB and HACD4 in the snow leopard (*Panthera uncia*; **Figure 5B**) and southern two-toed sloth (*Choloepus didactylus*; **Figure S7**). NCBI database mining reveals complete IFNN in additional Felidae and Phocidae species (**Figure 7B**), though it is pseudogenised in primates (Krause et al., 2005).

**Figure 7.**
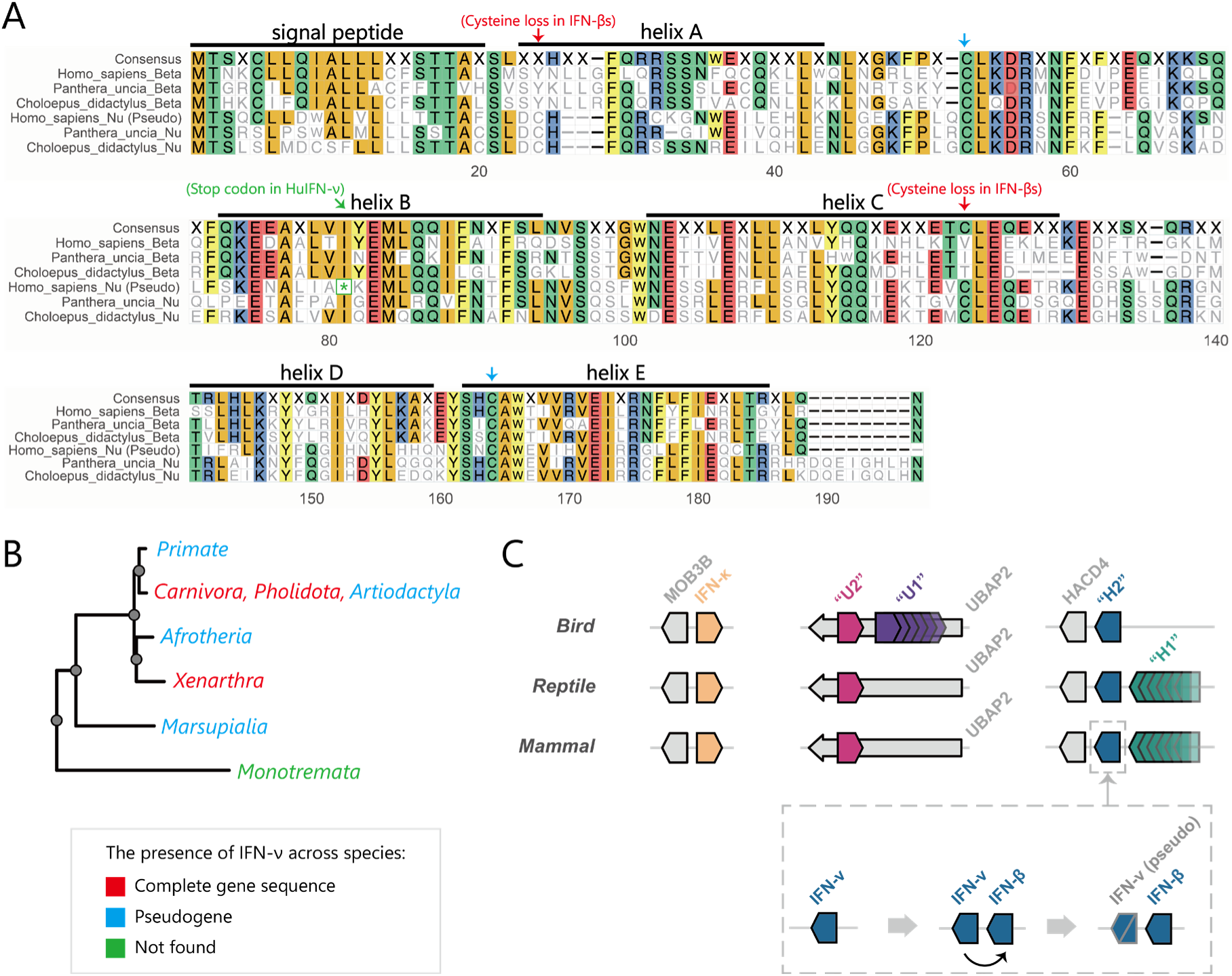
IFN-ν is the ancestor of mammalian IFN-Hs. **(A)** The MSA result of both IFN-ν and IFN-β in human (*Homo sapiens*), snow leopard (*Panthera uncia*), and southern two-toed sloth (*Choloepus didactylus*). Conserved cysteines are identified as blue arrow, while cysteine losses in IFN-β are identified as red arrow. **(B)** The presence of IFNN (encoding IFN-ν) in mammals. Gene status of “complete gene sequence (colored in red)”, “pseudogene (colored in blue)”, or “not found (colored in green)” are illustrated. **(C)** A general diagram of the relative patterns of most IFN-I genes in amniotes.

Based on above syntenic and phylogenetic analysis, we propose that 4C form is the ancestor of amnionate intronless IFN-I at MOB3B, UBAP2 and HACD4 loci, and IFN-υ may serve as the ancestor of mammalian IFN-Hs (**Figure 7C**).

### 8. Interchromosomal duplication of IFN-I genes

Intestingly, we identify 27 IFN-I genes from 7 species located outside the three conserved loci (MOB3B, HACD4, and UBAP2) **(Table S4 & Table S5)**. Among these, 21 IFN-I genes with adjacent genes show synteny with IFN-I flanking genes on human chromosome 9 **(Figure S6 & Table S4)**, strongly suggesting that chromosomal rearrangements contributed to their non-canonical genomic distribution. An additional five IFN-I genes are identified in Goode’s thornscrub tortoise (*Gopherus evgoodei*), distributed across chromosomes 1, 2, 3, and 24. In contrast, the canonical IFN-I loci (HACD4, MOB3B, and UBAP2) are located on chromosome 6 in this species. (**Figure 8A & Table S5)**. Notably, The tortoise chromosome 2 IFN-I flanked by MTRR and SEMA5A genes whose orthologs reside on disparate chromosomes in humans (chromosome 5), unstriped caecilian (chromosome 1), and coelacanth (chromosome 2) (**Figure 8B**). In contrast, the HACD4, MOB3B, and UBAP2 IFN-I loci are located on chromosome 9 (humans), chromosome 2 (caecilians), and chromosome 1 (coelacanths), respectively (**Figure 2E**). This syntenic inconsistency strongly supports an interchromosomal mechanism for this IFN-I gene duplication. Furthermore, the chromosome 2 IFN-I gene has also been identified in the Bolson tortoise (*Gopherus flavomarginatus*), providing additional support for the above conclusion (**Figure 8B**). Similarly, orthologs of the MTIF2 and CCDC88A genes flanking the tortoise chromosome 3 IFN-I gene are mapped to human chromosome 2, unstriped caecilian chromosome 3, and coelacanth chromosome 3, suggesting another independent interchromosomal duplication event (**Figure 8C**). Orthologs of the genes flanking the chromosome 1 IFN-I genes and chromosome 24 IFN-I gene are not located at the same chromosome, or even not identified, in human, caecilian, or coelacanth genomes, precluding subsequent syntenic analysis **(Table S5)**.

**Figure 8.**
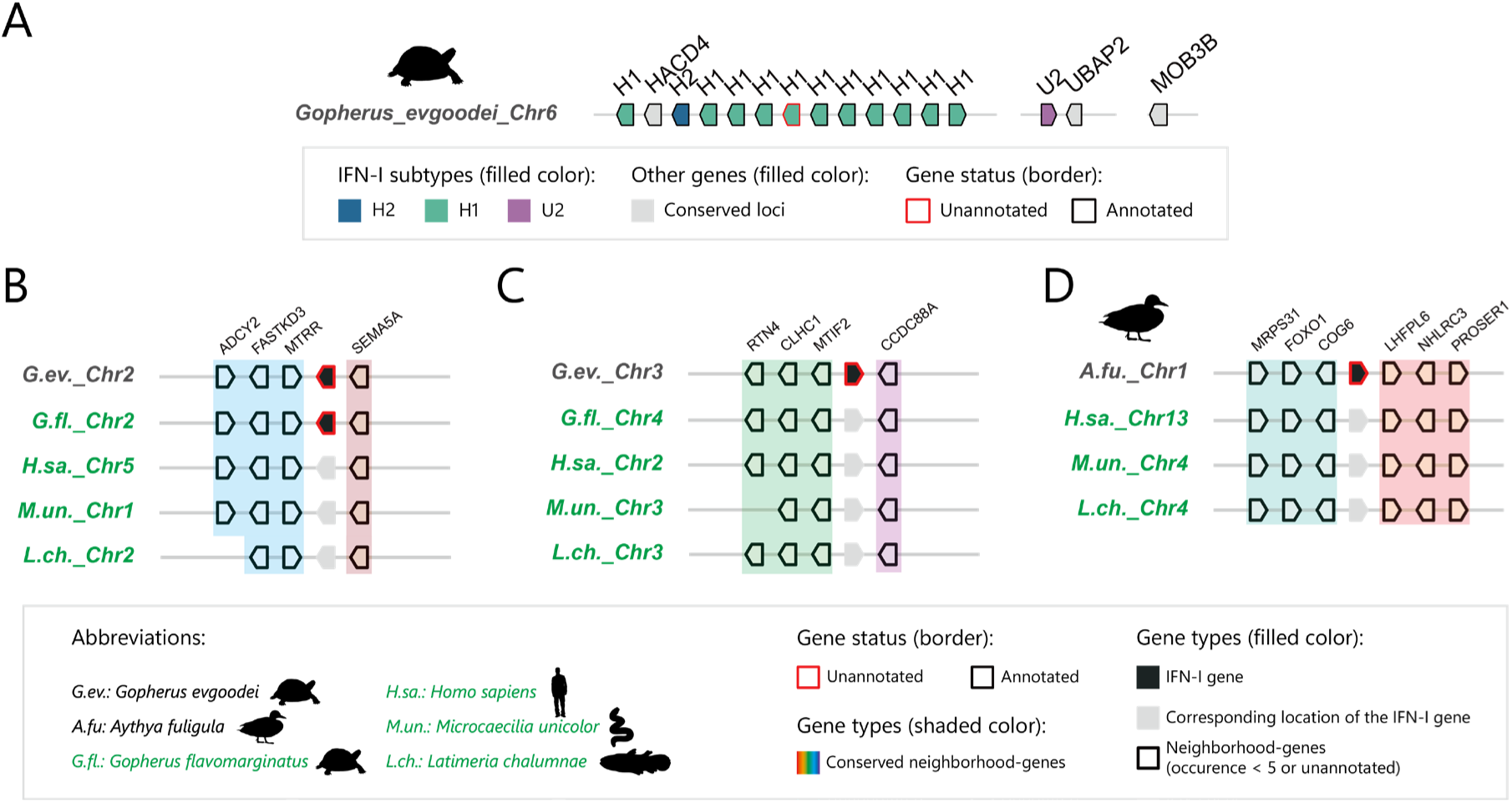
The illustration interchromosomal duplication of IFN-I genes in candidate species. **(A)** Canonical IFN-I distribution on Goode’s thornscrub tortoise (*Gopherus evgoodei*) chromosome 6. Gene arrows of IFN-I genes and corresponding conserved loci are differently colored. **(B)** Interchromosomal gene duplication events on the chromosome 2 of Goode’s thornscrub tortoise. IFN-I genes (colored in black) that are not flanked by common neighborhood-genes are presented, and conserved neighborhood-gene in Bolson tortoise (*Gopherus flavomarginatus* or *G.fl.*), human (*Homo sapiens* or *H.sa.*), unstriped caecilian (*Microcaecilia unicolor* or *M.un.*), and coelacanth (*Latimeria chalumnae* or *L.ch.*) are marked by different color shadings. **(C)** Interchromosomal gene duplication events on the chromosome 3 of Goode’s thornscrub tortoise. **(D)**. Interchromosomal gene duplication events on the chromosome 1 of tufted duck (*Aythya fuligula*). Only the first and the last IFN-I gene on the locus are presented in **(B)**, **(C)**, and **(D)**, and the gene orientations are adjusted to align the gene orientations in Goode’s thornscrub tortoise in these Figure.

While avian IFN-I genes are typically restricted to chromosome Z, we identify an exceptional case in tufted duck (*Aythya fuligula*), where one IFN-I gene (assigned as IFN-U2) is localized to chromosome 1, flanked by COG6 and LHFPL6 (thus referred to as “IFN-CL”, **Figure 8D**). Notably, the orthologs of these flanking genes show conserved synteny across distant vertebrates: human chromosome 13, caecilian chromosome 4, and coelacanth chromosome 4 (**Figure 8D**). Furthermore, IFN-I genes are found at the same locus in three additional diving duck species (*Aythya spp.*), but are absent in both the mallard *(Anas platyrhynchos*) and ruddy duck (*Oxyura jamaicensis*) (**Figure 9A**). These findings strongly support that an interchromosomal duplication of IFN-I genes occurred on chromosome 1 during the early divergence of diving ducks.

**Figure 9.**
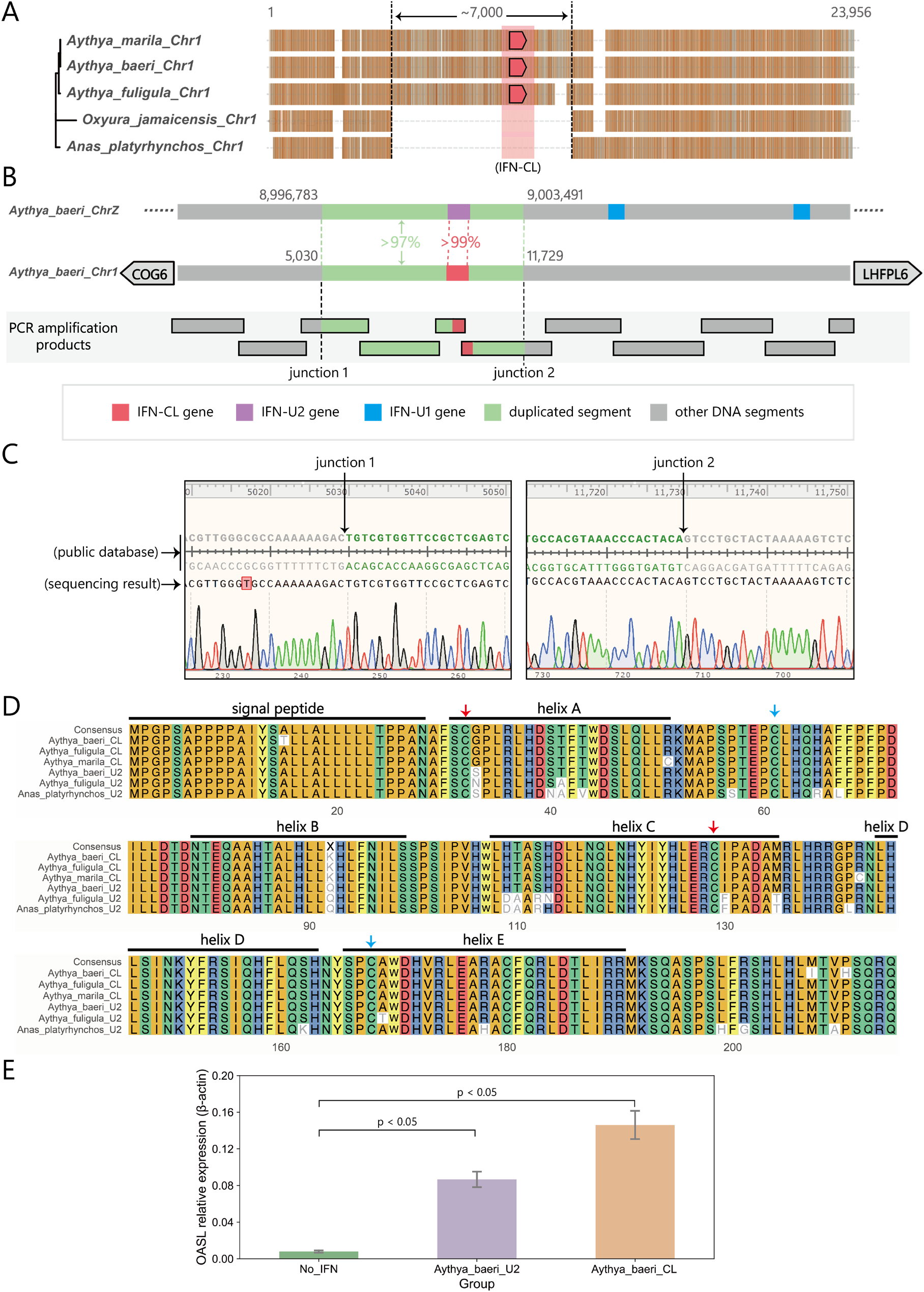
Unique IFN-CL gene is identified functional on the chromosome 1 of diving ducks. **(A)** The MSA results of DNA segment that from COG6 to LHFPL6 in diving ducks (*Aythya spp.*), mallard (*Anas platyrhynchos*) and ruddy duck (*Oxyura jamaicensis*) in public database. The numbers above represent the corresponding positions within the segment. **(B)** The results from PCR amplification and sequencing prove the existence of an IFN-CL gene in baer’s pochard (*Aythya baeri*). IFN-CL gene (red), IFN-U2 gene (purple), IFN-U1 gene (blue), duplicated segment (green), and other DNA segments (gray) are respectively colored. Corresponding PCR fragments are presented below. **(C)** Both nucleotide junctions of inserted segment in chromosome 1 in public database and sequencing results. **(D)** The MSA result of IFN-CL and IFN-U2 in ducks. IFN-U2 of *Aythya marila* is not found due to its low-quality genome assembly. **(E)** The induction of the OASL expression by IFN-CL and IFN-U2 in blood cells from baer’s pochard.

An approximately 7 kb DNA segment inserted at this locus shows high sequence homology to the region flanking the IFN-U2 gene on chromosome Z (99% identity for IFN-I coding region; 97% identity for whole segment) (**Figure 9B**). To confirm this insertion event, we amplify 11 DNA fragments by overlapping PCR using blood samples from Baer’s pochard (*Aythya baeri)* and sequenced the PCR products **(Materials & Methods 4.1)**. Both insertion boundaries (5’ and 3’ junctions) are unambiguously mapped, conclusively validating the insertion event (**Figure 9C & Figure S9)**. Consistent with its genomic origin, the IFN-CL protein sequences show remarkable similarity to diving duck IFN-U2s (**Figure 9D**).

To functionally characterize IFN-CL, we clone the coding sequences of Baer’s pochard IFN-CL and IFN-U2 into recombinant plasmids and express them in HEK293F cells. The supernatants are then used to stimulate peripheral blood mononuclear cells (PBMCs) **(Materials & Methods 4.2)**. Both IFN-CL and IFN-U2 significantly upregulate the expression of OASL (**Figure 9E**), a known interferon-stimulated gene (ISG) that is specifically induced by type I interferons across multiple species, including ducks **(Materials & Methods 4.3)**. These results demonstrate that IFN-CL is a functional type I interferon originating from an interchromosomal duplication event to diving duck chromosome 1.

## Discussion

It has been hypothesized that evolutionary retroposition event of intron-containing type I IFN transcripts gave arise to ‘intronless’ type I IFNs in amniotes during the period when vertebrates adapt the living environment from water to land^10–13^. To investigate this evolutionary of amniote IFN-I genes, we develop an integrated analytical pipeline combining the IFN-SCOPE classification model and GENE-GRADE algorithm. The computation results identify conserved IFN-I genes at three loci (HACD4, MOB3B, and UBAP2) across all major amniote lineages, leading us to propose that these three IFN-I loci were present in the most recent common ancestor (MRCA) of modern amniotes, and each locus evolved independently. Using synteny-guided phylogenetic analysis that considers chromosomal localization relative to conserved adjacent genes, position within IFN-I clusters and transcriptional orientation, we reveal the following findings: 1) IFNK orthologs are evolutionarily conserved across all major amniote lineages; 2) Convergent diversification patterns at HACD4 and UBAP2 loci produce single-copy IFN-H2/IFN-U2 variants and multicopy IFN-H1/IFN-U1 expansions; 3) The HACD4-proximal IFN-I gene (IFN-H2 subcluster) represents the ancestral form of this locus; 4) In birds with UBAP2-locus expansions, IFN-I gene proximal to the 3’-end of UBAP2 shows ancestral characteristics.

IFN-β exhibits unique characteristics among mammalian type I interferons, distinguished by both its induction mechanisms and functional properties^3,45,46^. While phylogenetic analyses initially proposed IFN-β as the ancestral mammalian IFN-I at HACD4 locus^9^, our synteny-guided evolutionary reconstruction identifies HACD4-proximal IFN-ν, with two evolutionaly conserved disulfide bonds, as the authentic progenitor of mammalian IFN-H (**Figure 7**). The evolutionary pattern of mammals at HACD4 locus, characterized by pervasive IFN-ν loss alongside conserved single-copy IFN-β, indicates functional redundancy between these IFN-H2 subtypes. This strongly suggests that precise dosage regulation is critical for their immunological functions. Notably, intact IFNN genes (encoding IFN-ν) persist in some carnivores (**Figure 7B**), while IFN-β shows lineage-specific expansions in ruminants (buffalo, sheep, goats, deer)^47,48^ and selected rodents^9^. These exceptions highlight the need to investigate the mechanisms driving unconventional IFN-H2 multiplication in specific lineages.

Our synteny-guided approach resolves longstanding ambiguity in avian IFN-I evolution: (1) confirming MOB3B-adjacent genes as genuine IFN-κ orthologs, and (2) establishing the single HACD4-locus gene as the true avian counterpart to mammalian IFN-H2 (IFN-ν/β/ε). This reclassification (**Figure 2, Figure 4, & Figure 5**) corrects prior interpretations^40,41^ and provides a validated evolutionary framework for comparative immunological studies.

Chicken IFN-U2 (chIFN-β) and IFN-U1 (chIFN-α) exhibit distinct biological activities and are considered evolutionary paralogs of their human counterparts (Sick et al., 1996; Schultz et al., 2004). Strikingly, comparative genomic analysis reveals noticable conservation of cis-regulatory elements in the IFN-U2 promoter across vertebrate lineages including mammalian IFN-β and selected IFN-I subtypes in amphibians and fishes^13,43^. This exceptional evolutionary preservation suggests strong selective pressure maintaining the transcriptional regulation of IFN-β-like genes. While the functional dichotomy between human IFN-β and IFN-α has been extensively characterized, with their distinct biological outcomes largely attributed to differential receptor binding affinities^49^. However, the evolutionary and functional significance of IFN-I diversification in non-mammalian amniotes represents a critical knowledge gap in comparative immunology.

Previous studies established local gene duplication as a major mechanism of IFN-I diversification in amniotes^8,9^. Our work extends this understanding by identifying interchromosomal duplication events in Goode’s thornscrub tortoise and tufted duck (**Figure 8**). Notably, we discovered an interchromosomal duplication on chromosome 1 during early diving duck radiation that generated IFN-CL (**Figure 9E**). Notably, IFN-CL represents an interchromosomal duplication of IFN-U2, which remains single-copy in most avian species. This evolutionary innovation in diving ducks holds particular significance given that wild and domestic ducks serve as key reservoirs for pandemic influenza viruses^50^, and diffent susceptibility exist between diving ducks and dabbling ducks (e.g. mallards)^51^. Thus, how this unconventional duplicaiotn of IFN-CL contributed to susceptibility to influenza or other viral infection worth to be investigated in the future.

In summary, these findings significantly advance our comprehension of amniote IFN-I evolution by demonstrating locus-specific diversification, while providing a framework for predicting IFN-I functions through comparative immunology. Moreover, this work provides an evolutionary blueprint for developing novo IFN-I based antiviral therapies.

## Authors’ Contributions

**Le Zhang:** Writing – review & editing, Supervision, Project administration, Funding acquisition. **Fubo Ma:** Conceptualization, Data preprocessing, Data computation, Data analysis, Investigation, Data curation, Visualization, Writing – original draft. **Jinpeng Liu:** Provide samples of the baer’s pochard (*Aythya baeri*). **Yangchao Yu:** Data preprocessing, Data computation, Data analysis, Data curation. **Junxiao Ma:** Conduct IFN-I experiments using the samples. **Lei Zhang:** Data preprocessing, Data computation, Data analysis, Data curation. **Bing Li:** Writing – review & editing. **Chaofan Li:** Conduct IFN-I experiments using the samples. **Peng Liu:** Writing – review & editing, Supervision. **Liguo Zhang:** Conceptualization, Data analysis, Investigation, Writing – review & editing, Supervision, Project administration, Funding acquisition. All authors have read and approved the final manuscript.

## Acknowledgements

This work was supported by grants from National Science and Technology Major Project (2021YFF1201200 and 2024ZD0532900), National Natural Science Foundation of China (62372316), and Key Projects of Sichuan Provincial Department of Science and Technology (2024YFHZ0091 and 2025YFHZ0066).

