## Supplementary Information for "Locus-specific Convergent Evolution and Interchromosomal Rearrangements Contribute Type I Interferon Diversification in Amniotes"

### Supplementary Tables

**Table S1.** Data sources and computation results in this paper.

**Table S2.** PCR primers used in this paper.

**Table S3.** Amniote IFN-Is that cannot be accurately assigned into specific subtypes.

**Table S4.** IFN-I gene with neighborhood genes that are located on the chromosome 9 of human.

**Table. S5.** IFN-I gene with neighborhood genes that are NOT located on the chromosome 9 of human.

### Supplementary Figures

**Figure S1.** Overview of the phylogenetic relationships of all candidate species in this study.

**Figure S2.** The flowchart of the IFN-SCOPE model.

**Figure S3.** The flowchart of GENE-GRADE algorithm.

**Figure S4.** The distribution of all IFN-I genes at conserved loci or "exception" category in mammals.

**Figure S5.** The distribution of all IFN-I genes at conserved loci or "exception" category in birds and reptiles.

**Figure S6.** The presence of IFN-I neighborhood-genes that on conserved loci or "exception" category of our candidate species.

**Figure S7.** Proposed novel nomenclature of IFN-I in mammals of our candidate species.

**Figure S8.** Proposed novel nomenclature of IFN-I in birds and reptiles of our candidate species.

**Figure S9.** The comparisons and identities of DNA sequence in public database and by sequencing.

**Table S1. Data sources and computation results in this paper.**

Data sources include whole genome assemblies, gff files, IFN-I gene and protein sequence data (annotated IFN), and data version for candidate species; computation results include details of unannotated IFN-Is.

| Data type | Data name | Download/Analysis link |
|---|---|---|
| Data sources | Whole genome assemblies | https://www.ncbi.nlm.nih.gov/assembly/ |
| | gff files | https://www.ncbi.nlm.nih.gov/assembly/ |
| | Annotated IFN-Is | https://interferon.netlify.app/download |
| | Data version | https://interferon.netlify.app/dataVersion |
| Computation results | Unannotated IFN-Is | https://interferon.netlify.app/download |

**Table S2. PCR primers used in this paper.**

Because the genome of baer's pochard has not been annotated during experimentation and manuscript preparation, the primers of corresponding genes are designed individually based on BLAST results.

| Gene name | Forward (5' to 3') | Reverse (5' to 3') |
|---|---|---|
| IFN-CL | CCCCACCTTGTCGTTGTCCAGC | CCCAGCCAATCTATCTATCCAG |
| | ATCTTCAAACGCTTGGAGTTACAT | GCTTTATACCTGAGCACAGAGCC |
| | TTCCCAGCAAAGCCAATAACAT | GCAGTAGGAGAACAACGAGCAGT |
| | ATGACCAACCTTCTGGCGATGC | CCTCTGGATTCCCTTGTATGTG |
| | ACTCCCTTAGGTTCCATTGAAGA | CGAGGTGGTAGATGTAGTGGTTGA |
| | CCTCCTCAAACACCTCTTCAACATC | GTCCAGCAACCTGAACTGTAAACAC |
| | TTTCTGCCATGTTTCCTGTGAA | AGCATCAGCATTTGTTGCCTTC |
| | GCTTCTTTCTACTTCTGTCCCATAC | TCATAACTACCCTATTCCAGTCCTAA |
| | GCCAGGGAAGGCAGCCTCACAT | CACCTCCTCAGAGCAGAGCAATAACAAA |
| | GGCGTTTGATCCAGTCTGATTT | GATTTCCTCCTCCTGCTCCTTC |
| | CATGTTCCCGGACTGTAAAAGA | CAAAGCGACTGTCCCCTTGGTC |
| GAPDH | CGTGTGGTGGACTTGATGGT | GAGGAGTGGGGGAGACAGAA |
| OASL | AGAATCGGCTCCAAGAGTGC | ATCGTAGGTGGGCAGGATGT |

**Table S3. Amniote IFN-Is that cannot be accurately assigned into specific subtypes.**

Due to the limitations of the quality of genomic data, three amniote IFN-I genes cannot be accurately assigned since they do not group with any other IFN-I.

| Species | Conserved loci | Gene name |
|---|---|---|
| *Oxyura jamaicensis* | “exception” category | NW_023312189.1_POS988 |
| *Oxyura jamaicensis* | “exception” category | NW_023312179.1_POS7562 |
| *Rhinolophus ferrumequinum* | HACD4 | NC_046295.1_POS18678887 |

**Table S4. IFN-I gene with neighborhood genes that are located on the chromosome 9 of human.**

*H.sa.: Homo sapiens; M.un.: Microcaecilia unicolor; L.ch.: Latimeria chalumnae.*

| Species | Gene name | Chromosome | Gene category | Neighborhood-gene | | Neigh. gene on *H.sa.* chromosome | | Neigh. gene on *M.un.* chromosome | | Neigh. gene on *L.ch.* chromosome | |
|---|---|---|---|---|---|---|---|---|---|---|---|
| | | | | L | R | L | R | L | R | L | R |
| *Trichosurus vulpecula* | LOC118831614 | Chr9 | H1 | MTAP | SYK | Chr9 | Chr9 | Chr2 | Chr2 | Chr1 | Chr1 |
| *Trichosurus vulpecula* | LOC118831668 | Chr9 | H1 | MTAP | SYK | Chr9 | Chr9 | Chr2 | Chr2 | Chr1 | Chr1 |
| *Trichosurus vulpecula* | LOC118831665 | Chr9 | H1 | MTAP | SYK | Chr9 | Chr9 | Chr2 | Chr2 | Chr1 | Chr1 |
| *Trichosurus vulpecula* | LOC118831684 | Chr9 | H1 | MTAP | SYK | Chr9 | Chr9 | Chr2 | Chr2 | Chr1 | Chr1 |
| *Ornithorhynchus anatinus* | IFN4 | ChrX5 | H2 | TBC1D2 | TRIM41 | Chr9 | Chr5 | Chr2 | No gene | Chr1 | No gene |
| *Ornithorhynchus anatinus* | IFN3 | ChrX5 | H2 | TBC1D2 | TRIM41 | Chr9 | Chr5 | Chr2 | No gene | Chr1 | No gene |
| *Ornithorhynchus anatinus* | IFNK | ChrX5 | H2 | TBC1D2 | TRIM41 | Chr9 | Chr5 | Chr2 | No gene | Chr1 | No gene |
| *Ornithorhynchus anatinus* | LOC114807941 | ChrX5 | H2 | TBC1D2 | TRIM41 | Chr9 | Chr5 | Chr2 | No gene | Chr1 | No gene |
| *Ornithorhynchus anatinus* | IFN1 | ChrX5 | H2 | TBC1D2 | TRIM41 | Chr9 | Chr5 | Chr2 | No gene | Chr1 | No gene |
| *Ornithorhynchus anatinus* | LOC114807967 | ChrX5 | H2 | TBC1D2 | TRIM41 | Chr9 | Chr5 | Chr2 | No gene | Chr1 | No gene |
| *Tachyglossus aculeatus* | LOC119948206 | ChrX4 | H2 | TRIM41 | TBC1D2 | Chr5 | Chr9 | No gene | Chr2 | No gene | Chr1 |
| *Tachyglossus aculeatus* | LOC119948164 | ChrX4 | H2 | TRIM41 | TBC1D2 | Chr5 | Chr9 | No gene | Chr2 | No gene | Chr1 |
| *Tachyglossus aculeatus* | LOC119947822 | ChrX4 | H2 | TRIM41 | TBC1D2 | Chr5 | Chr9 | No gene | Chr2 | No gene | Chr1 |
| *Tachyglossus aculeatus* | LOC119947800 | ChrX4 | H2 | TRIM41 | TBC1D2 | Chr5 | Chr9 | No gene | Chr2 | No gene | Chr1 |
| *Mauremys reevesii* | LOC120407278 | Chr6 | H1 | SYK | MTAP | Chr9 | Chr9 | Chr2 | Chr2 | Chr1 | Chr1 |
| *Mauremys reevesii* | LOC120407297 | Chr6 | H1 | SYK | MTAP | Chr9 | Chr9 | Chr2 | Chr2 | Chr1 | Chr1 |
| *Mauremys reevesii* | LOC120407300 | Chr6 | H1 | SYK | MTAP | Chr9 | Chr9 | Chr2 | Chr2 | Chr1 | Chr1 |
| *Mauremys reevesii* | LOC120407541 | Chr6 | H1 | SYK | MTAP | Chr9 | Chr9 | Chr2 | Chr2 | Chr1 | Chr1 |
| *Mauremys reevesii* | LOC120407302 | Chr6 | H1 | SYK | MTAP | Chr9 | Chr9 | Chr2 | Chr2 | Chr1 | Chr1 |
| *Gopherus evgoodei* | LOC115653518 | Chr6 | H1 | MTAP | KIAA2026 | Chr9 | Chr9 | Chr2 | Chr2 | Chr1 | Chr1 |
| *Corvus moneduloides* | NC_045511.1_ POS26566131 | ChrZ | κ | CAST | C9ORF72 | Chr5 | Chr9 | Chr2 | Chr2 | Chr1 | Chr1 |

**Table. S5. IFN-I gene with neighborhood genes that are NOT located on the chromosome 9 of human.**

*H.sa.: Homo sapiens; M.un.: Microcaecilia unicolor; L.ch.: Latimeria chalumnae.*

| Species | Gene name | Chromosome | Gene category | Neighborhood gene | | Neigh. gene on *H.sa.* chromosome | | Neigh. gene on *M.un.* chromosome | | Neigh. gene on *L.ch.* chromosome | |
|---|---|---|---|---|---|---|---|---|---|---|---|
| | | | | L | R | L | R | L | R | L | R |
| *Gopherus evgoodei* | NC_044345.1_POS12357003 | Chr24 | H1 | CADM3 | No gene | Chr1 | No gene | Chr14 | No gene | Chr21 | No gene |
| *Gopherus evgoodei* | NC_044322.1_POS221191147 | Chr1 | H1 | EPHA1 | YBX3 | Chr7 | Chr12 | Chr14 | Chr14 | No gene | No gene |
| *Gopherus evgoodei* | NC_044322.1_POS221199478 | Chr1 | H1 | EPHA1 | YBX3 | Chr7 | Chr12 | Chr14 | Chr14 | No gene | No gene |
| *Gopherus evgoodei* | NC_044323.1_POS129589677 | Chr2 | H1 | MTRR | SEMA5A | Chr5 | Chr5 | Chr1 | Chr1 | Chr2 | Chr2 |
| *Gopherus evgoodei* | NC_044324.1_POS198858452 | Chr3 | H1 | MTIF2 | CCDC88A | Chr2 | Chr2 | Chr3 | Chr3 | Chr3 | Chr3 |
| *Aythya fuligula* | LOC116501035 | Chr1 | U2 | COG6 | LHFPL6 | Chr13 | Chr13 | Chr4 | Chr4 | Chr4 | Chr4 |

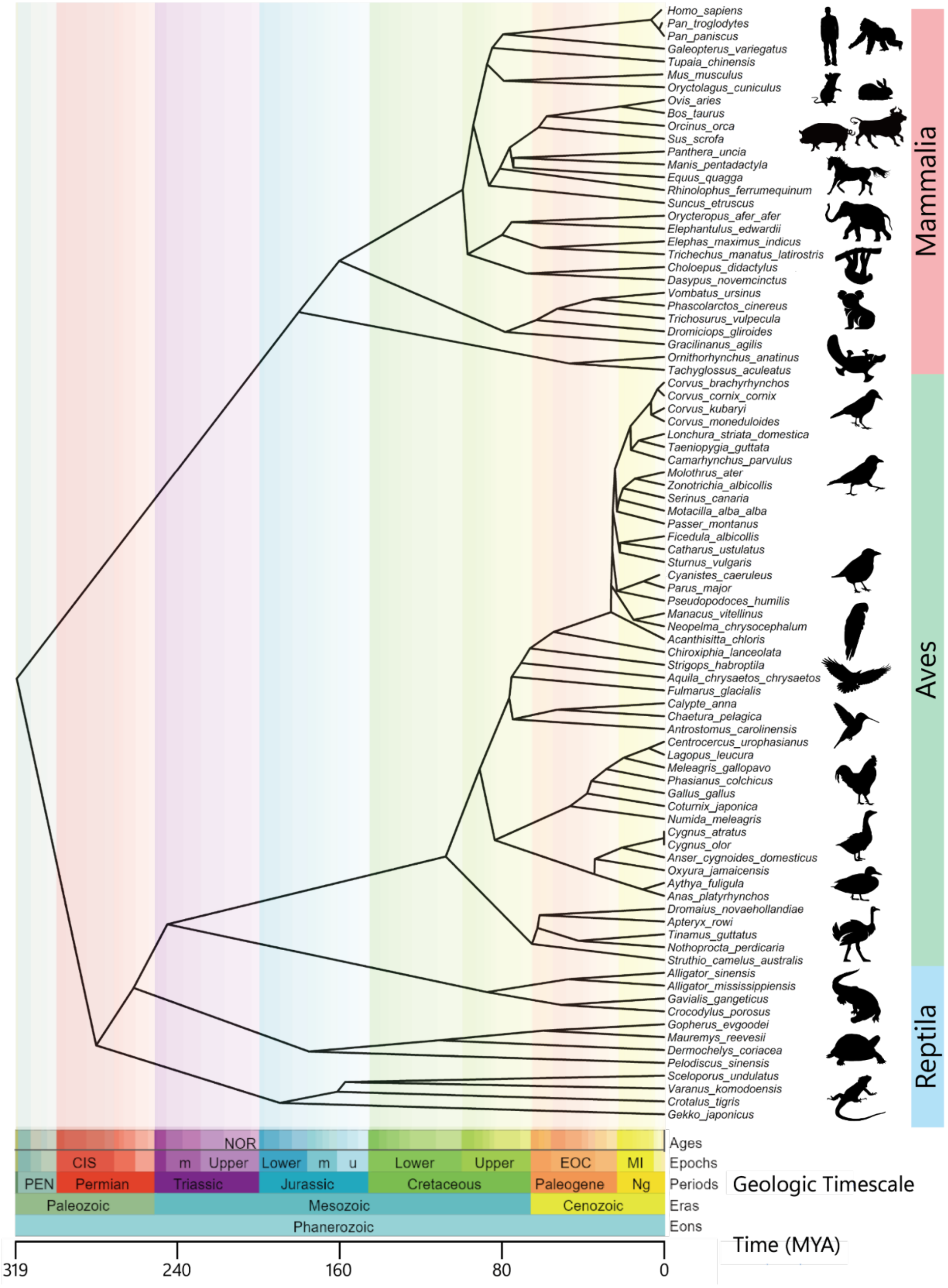


**Figure S1. Overview of the phylogenetic relationships of all candidate species in this study.**

According to the taxonomy of amniotes, candidate mammals (Mammalia, above), birds (Aves, middle) and reptiles (Reptilia, below) are presented in the Figure. The geological timescales and estimated divergence times are provided at the bottom of the Figure. The Figure is curated from TimeTree (Kumar, et al., 2022) and silhouette icons are obtained from Flaticon.

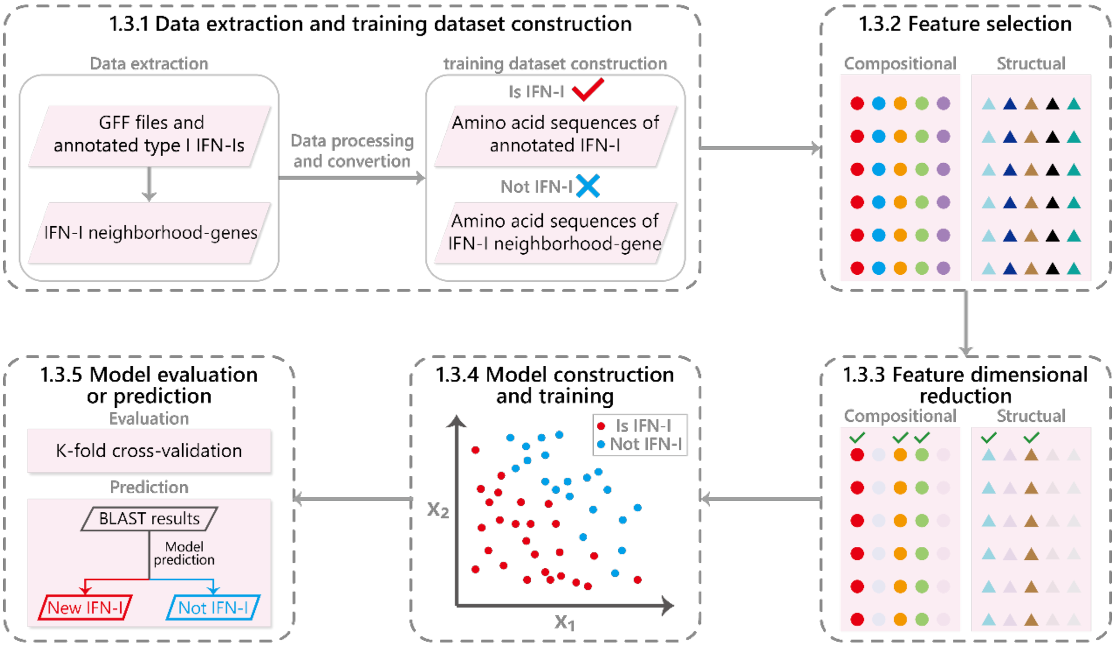


**Figure S2. The flowchart of the IFN-SCOPE model.**

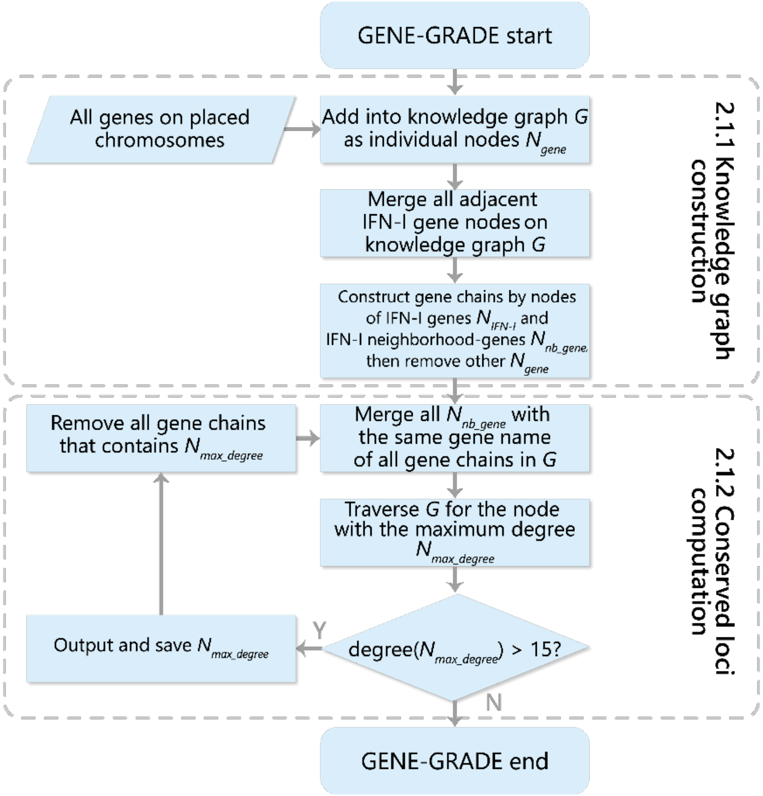


**Figure S3. The flowchart of GENE-GRADE algorithm.**

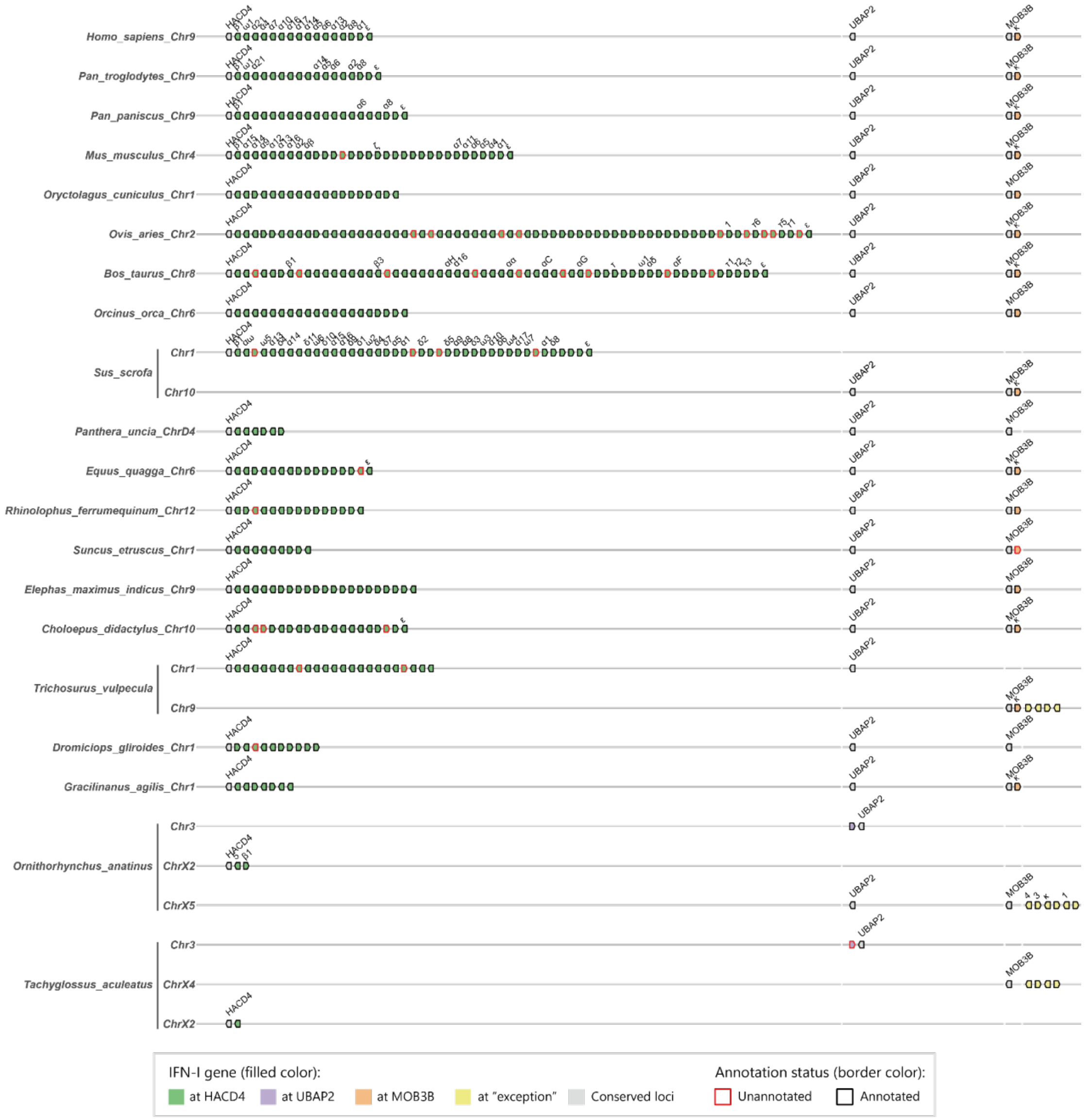


**Figure S4. The distribution of all IFN-I genes at conserved loci or "exception" category in mammals.**

IFN-I genes on the chromosomes (letter or number after the scientific name) of mammals are displayed according to their corresponding conserved loci.

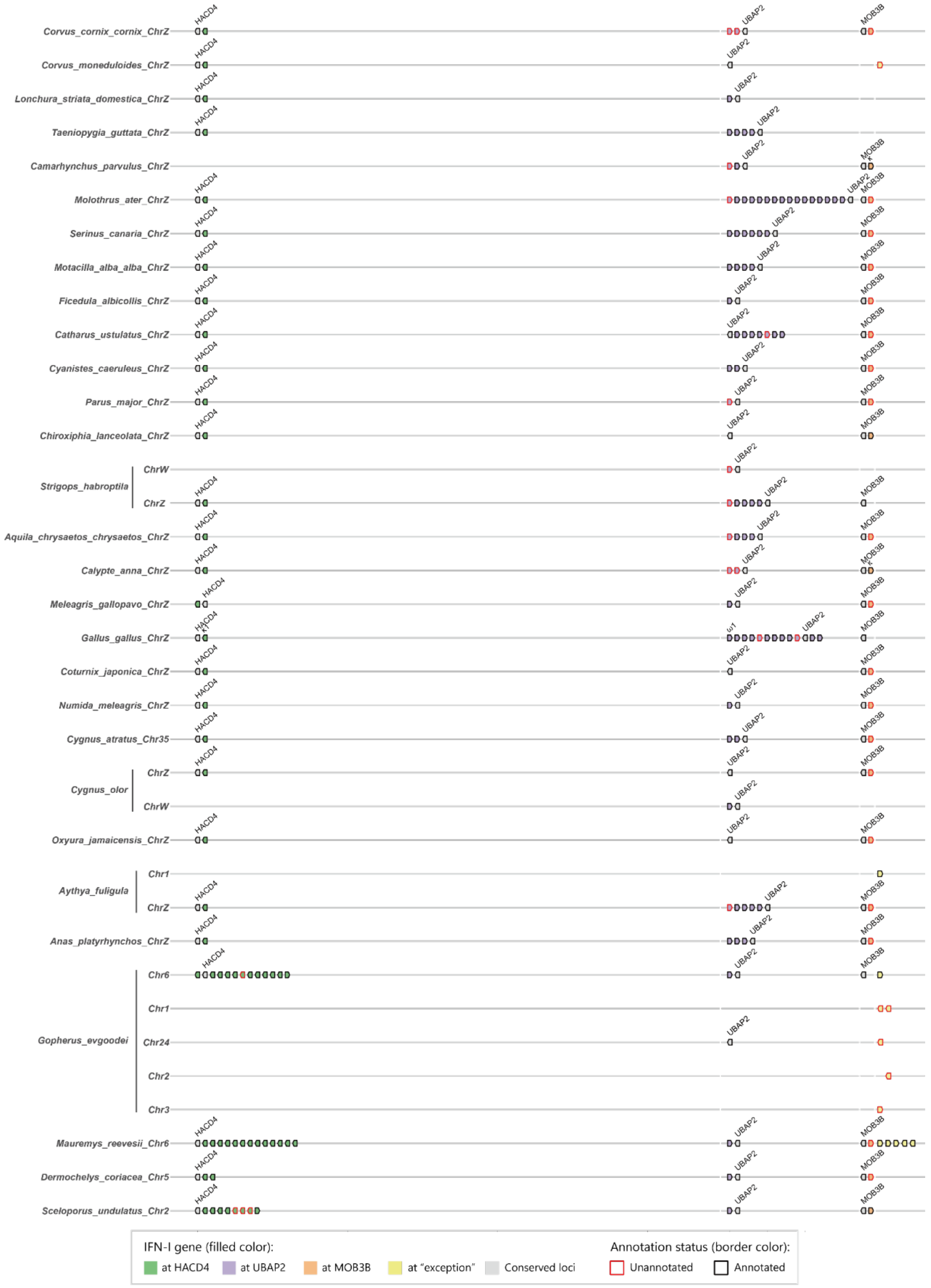


**Figure S5. The distribution of all IFN-I genes at conserved loci or "exception" category in birds and reptiles.**

IFN-I genes on the chromosomes (letter or number after the scientific name) of birds and reptiles are displayed according to their corresponding conserved loci.

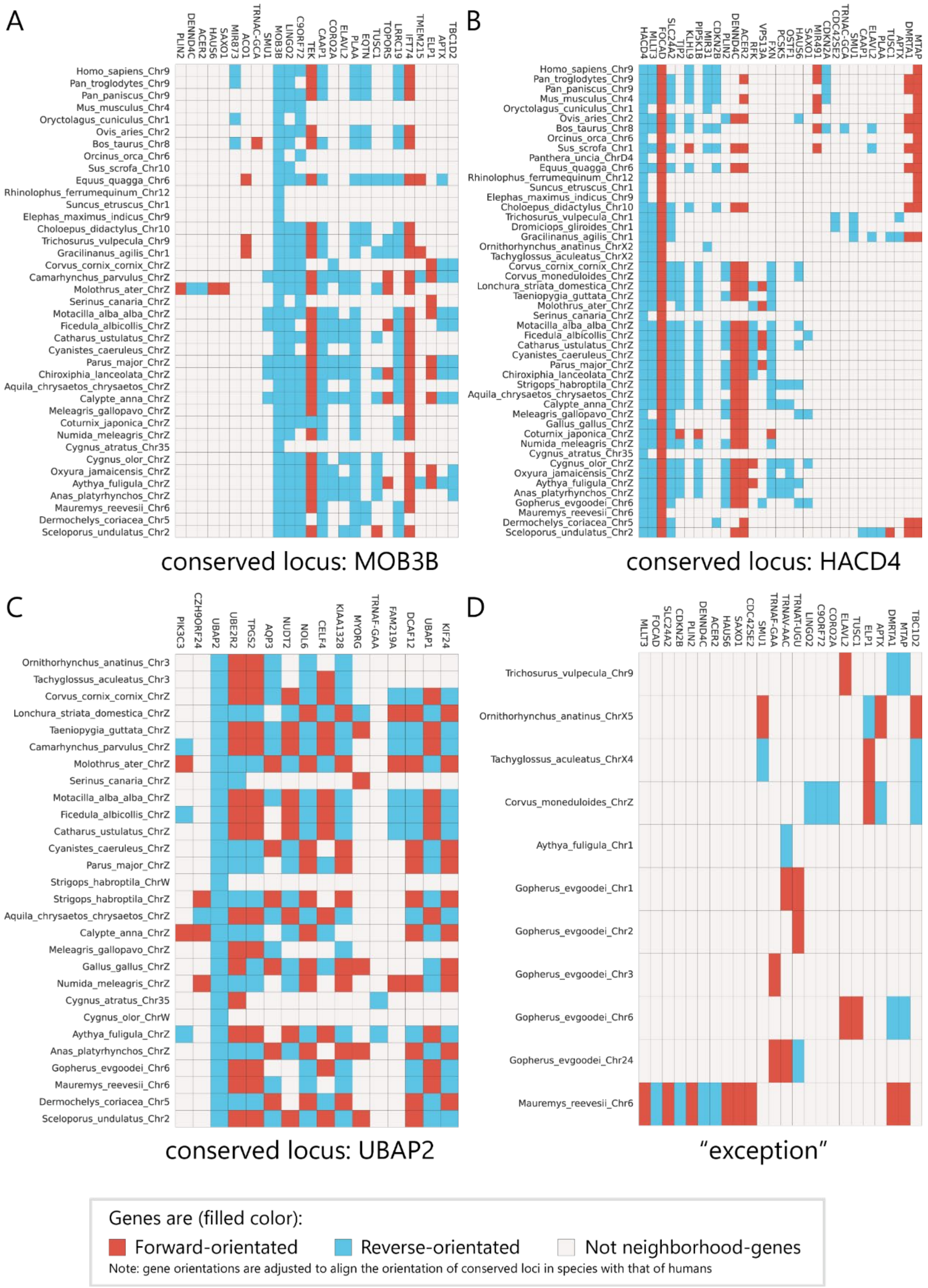


**Figure S6. The presence of IFN-I neighborhood-genes that on conserved loci or "exception" category in our candidate species.**

In this Figure, neighborhood-gene of IFN-Is (the X-axis) at MOB3B locus **(A)**, HACD4 locus **(B)**, UBAP2 locus **(C)**, and "exception" category **(D)** of chromosomes (the Y-axis) are accordingly displayed and colored by their gene orientation in each subplot. Only neighborhood-genes occurring more than five times are presented in the figure.

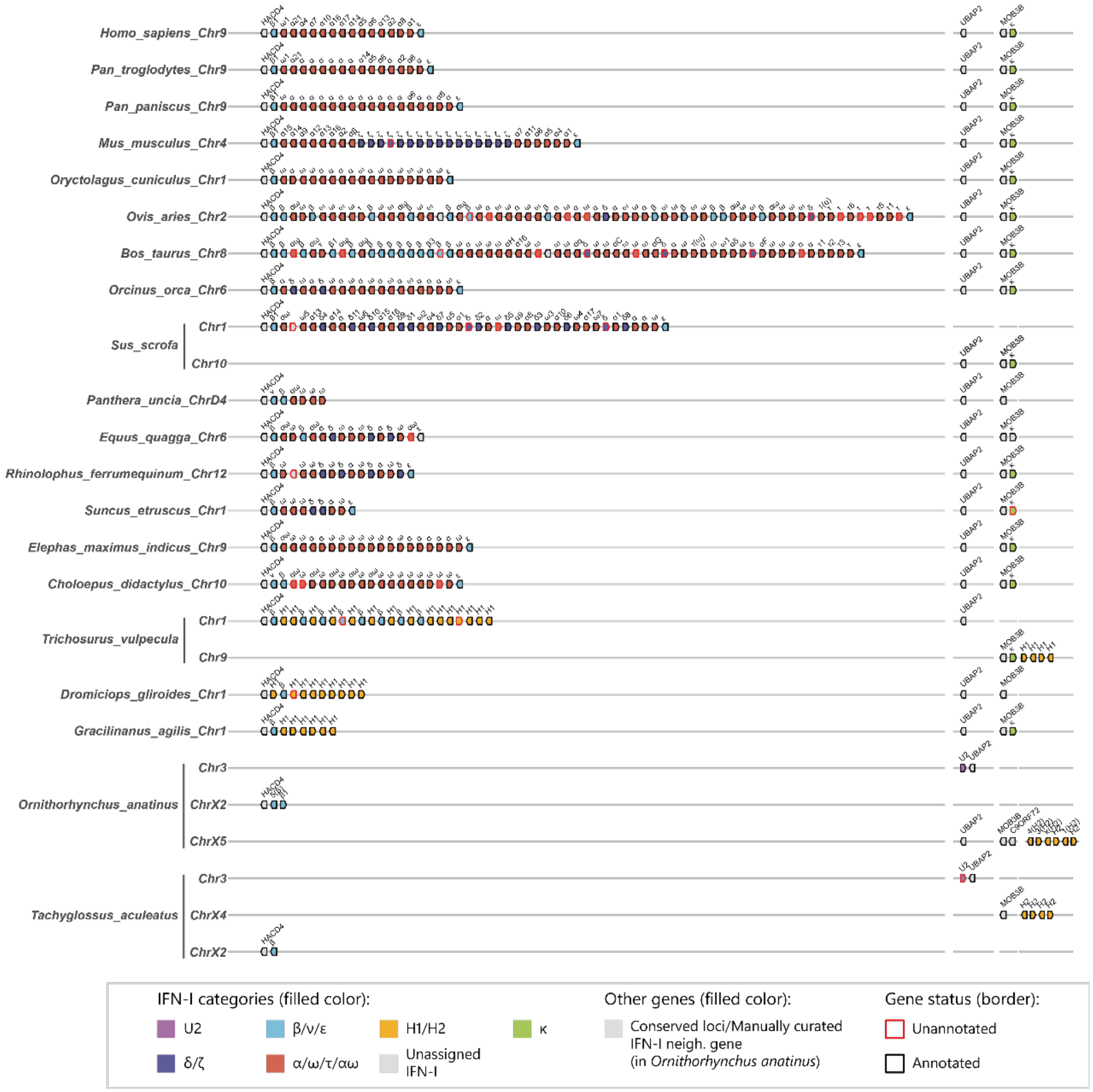


**Figure S7. Proposed novel nomenclature of IFN-I in mammals of our candidate species.** The color of each IFN-I gene indicates the proposed novel nomenclature in this study. We have highlighted C9ORF72 in platypus separately for illustrative purpose in the main text.

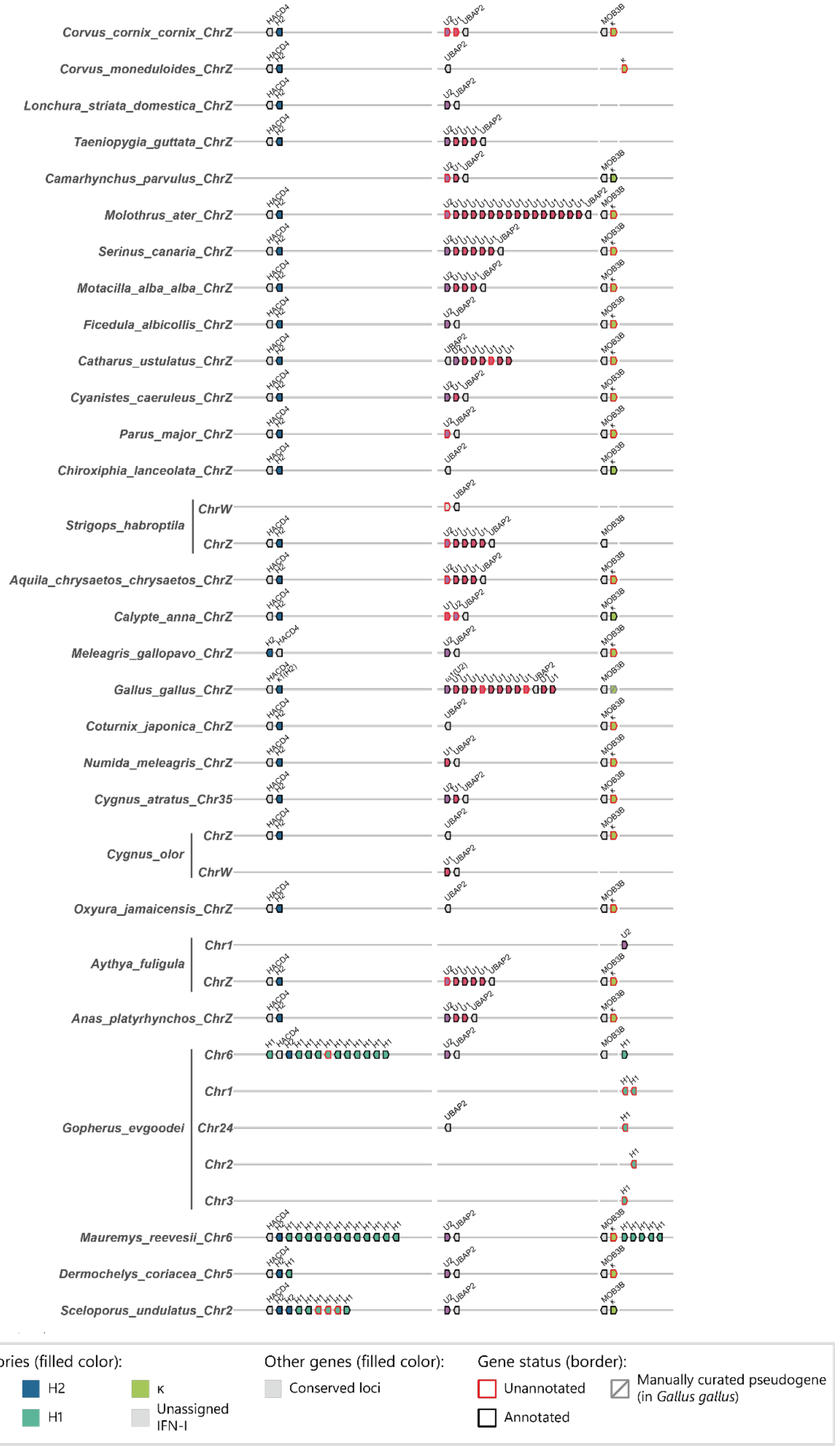

Corvus_cornix_cornix_ChrZ
Corvus_moneduloides_ChrZ
Lonchura_striata_domestica_ChrZ
Taeniopygia_guttata_ChrZ
Camarhynchus_parvulus_ChrZ
Molothrus_ater_ChrZ
Serinus_canaria_ChrZ
Motacilla_alba_alba_ChrZ
Ficedula_albicollis_ChrZ
Catharus_ustulatus_ChrZ
Cyanistes_caeruleus_ChrZ
Parus_major_ChrZ
Chiroxiphia_lanceolata_ChrZ
Strigops_habroptila
ChrW
ChrZ
Aquila_chrysaetos_chrysaetos_ChrZ
Calypte_anna_ChrZ
Meleagris_gallopavo_ChrZ
Gallus_gallus_ChrZ
Coturnix_japonica_ChrZ
Numida_meleagris_ChrZ
Cygnus_atratus_Chr35
Cygnus_olor
ChrZ
ChrW
Oxyura_jamaicensis_ChrZ
Aythya_fuligula
Chr1
ChrZ
Anas_platyrhynchos_ChrZ
Gopherus_evgoodei
Chr6
Chr1
Chr24
Chr2
Chr3
Mauremys_reevesii_Chr6
Dermochelys_coriacea_Chr5
Sceloporus_undulatus_Chr2
HACD4
UBAP2
MOB3B
IFN-I categories (filled color):
U2
U1
H2
H1
κ
Unassigned IFN-I
Other genes (filled color):
Conserved loci
Gene status (border):
Unannotated
Annotated
Manually curated pseudogene (in Gallus gallus)

**Figure S8. Proposed novel nomenclature of IFN-I in birds and reptiles of our candidate species.**

The color of each IFN-I gene indicates the proposed novel nomenclature in this study. We have highlighted the pseudo gene of IFN-κ in chicken (*Gallus gallus*) separately for illustrative purpose in the main text.

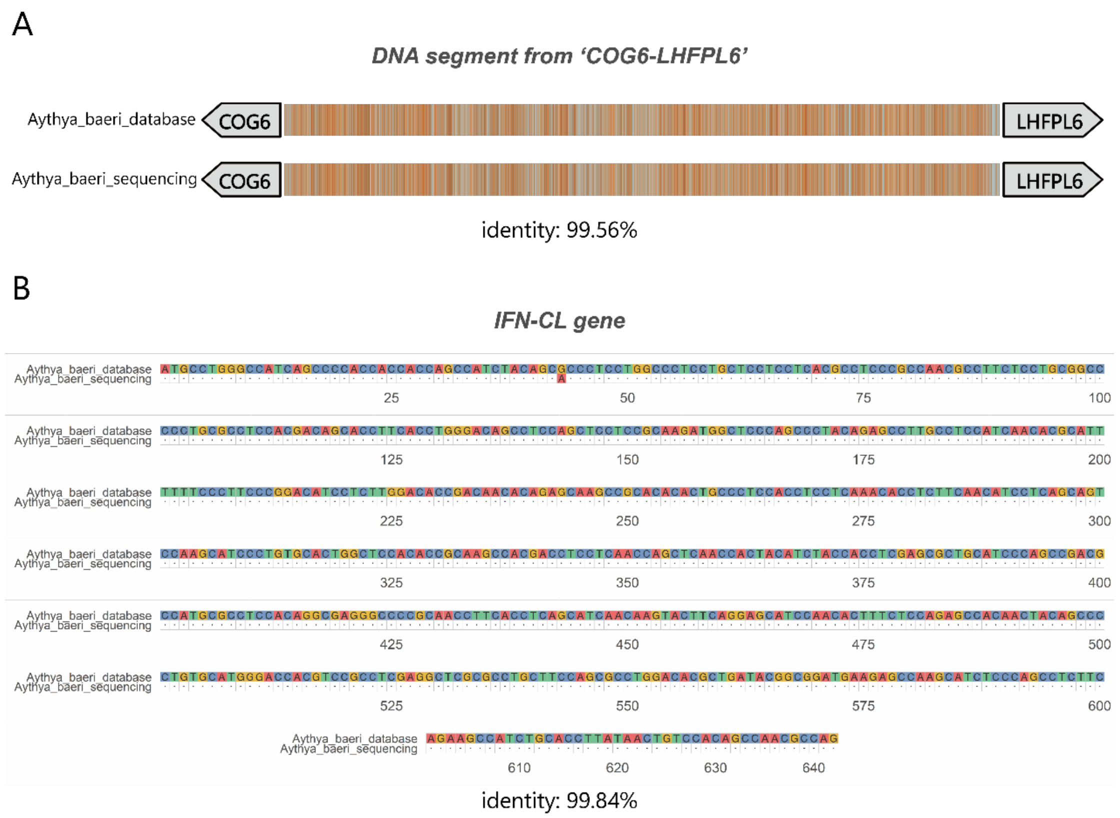


**Figure S9. The comparisons and identities of DNA sequence in public database and by sequencing.**

**(A)** The DNA segment that from COG6 to LHFPL6. **(B)** The IFN-CL gene.
